# It’s Not Rewarding for Mitochondria: Dopamine-Induced Mitochondrial Dysfunction Activates cGAS-STING to Drive IL-6 Secretion in Macrophages

**DOI:** 10.64898/2026.04.23.719926

**Authors:** Marzieh Daniali, Breana Channer, Erin O. Curley, Amirali Amirfallah, Taylor Kist, John Montilla, Stephanie Kosashvili, Lexi Sheldon, Kelly Stauch, Joshua G. Jackson, Anneliese M. Faustino, Thomas Beer, Hsin-Yao Tang, Stephanie M. Matt, Will Dampier, Howard Fox, Peter J. Gaskill

## Abstract

Despite increasing data demonstrating dopamine as an inflammatory mediator of the innate immune system, the molecular mechanisms underlying its effects in human cells remain incompletely defined. Here, we define an unrecognized pathway in which dopamine induces robust IL-6 secretion in primary human monocyte-derived macrophages (hMDMs) through mitochondrial stress. Dopamine initiates a transient mitochondrial membrane depolarization that leads to sustained alterations in mitochondrial dynamics, including morphology and metabolism, in a time-dependent manner. These events promote the mtDNA release into the cytoplasm, triggering cGAS-STING pathway and downstream NF-κB signaling. Pharmacological inhibition at multiple nodes of this pathway attenuates IL-6 secretion, establishing mitochondrial dysfunction and cGAS-STING signaling as central mediators of dopamine-driven IL6 secretion. Variability in dopamine receptor expression across donors correlates with the magnitude of IL-6 responses. Together, these findings redefine the interface between dopamine signaling and systemic inflammation and highlight an unrecognized source of inter-individual variation in immune responses.

**Graphical Abstract:** 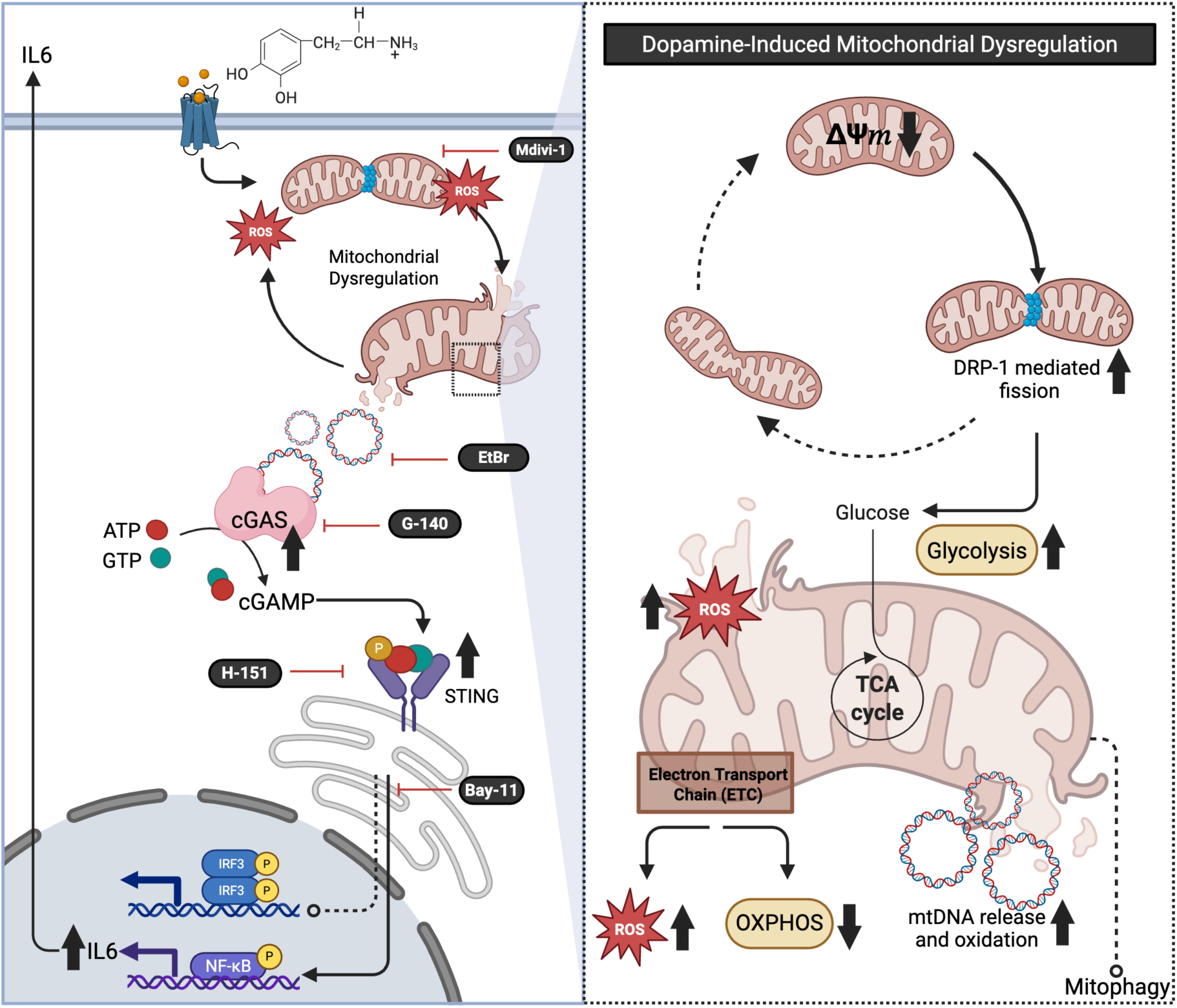

Dopamine induces mitochondrial dysfunction mediated through dopamine receptors’ signaling. This includes alterations in mitochondrial membrane potential, leading to excessive DRP1-mediated mitochondrial fission, increased production of mitochondrial superoxide, and metabolic reprogramming toward enhanced glycolysis with reduced oxidative phosphorylation. Sustained mitochondrial damage is further exacerbated by impaired mitophagy, resulting in the release of mitochondrial DNA (mtDNA) into the cytoplasm. Cytosolic mtDNA, acting as a double-stranded DNA ligand, activates the cGAS-STING pathway, which subsequently induces NF-κB signaling, ultimately driving the production and secretion of the pro-inflammatory cytokine IL-6. Created on Biorender.com.

## Introduction

Dopamine is a catecholamine neurotransmitter that is widely studied for its role in reward processing, movement, and cognition^1,2^. Beyond its classical role in neurotransmission, an increasing number of studies show that dopamine also mediates bidirectional signaling between the nervous and immune systems, acting as an immunomodulatory factor^3–5^. Relatively high concentrations of dopamine are present in both the central nervous system and peripheral organs, and a broad range of immune cells express dopamine receptors, providing the infrastructure for dopamine to regulate the immune system throughout the body^4,6,7^. Dopamine concentrations vary between organs and different physiological and pathological conditions can induce varying levels of dopamine concentrations, and therefore the magnitude of dopamine-driven immune responses may differ^4,8^.

The precise mechanism(s) by which dopamine drives these processes is not clear, but elevated dopamine levels can modulate inflammatory responses in a variety of immune cells, including myeloid cells. In human cells, these effects are primarily pro-inflammatory^4,9–12^. In contrast, studies in rodent models report both pro- and anti-inflammatory effects of dopamine, with more anti-inflammatory findings, highlighting a species-specific divergence in dopaminergic immunomodulation^4,13,14^. These differences are thought to arise, at least in part, from distinct dopamine receptor expression patterns^9,12^. Human myeloid cells express both D1-like (DRD1 and DRD5) and D2-like (DRD2, DRD3, and DRD4) receptors, with expression levels varying across cell types (e.g., macrophages versus microglia) and across the population^9,15^. In myeloid cells, while all dopamine receptors are expressed, D1-like receptors are generally expressed at higher levels than D2-like receptors^4,16^. These expression levels may be relevant to regulatory function, as a recent study showed dopamine-induced IL-1β production in human macrophages and iPSC-derived microglia correlates with the relative expression of dopamine receptor subtypes^9^. This suggests receptor balance may be an important determinant of dopaminergic inflammatory signaling, although the same correlations have not been shown for the secretion of other cytokines in human myeloid cells.

Nonetheless, dopamine does induce other pro-inflammatory cytokines in human myeloid cells, implicating this catecholamine as an important driver of innate immune activity^17,18^. Among the cytokines influenced by dopamine, interleukin-6 (IL-6) is a key mediator linking innate and adaptive immune responses. IL-6 is produced by multiple immune cell types, including myeloid cells as well as T and B lymphocytes, and plays essential roles in host defense, tissue repair, and immune regulation^19,20^. Dysregulated IL-6 production contributes to chronic inflammatory and autoimmune diseases, such as Rheumatoid Arthritis (RA), Systemic Lupus Erythematosus (SLE), and Inflammatory Bowel Disease (IBD). Changes in IL-6 are also strongly linked to depression, anxiety and stress-disorders, with the disruption of CNS dopamine metabolism connecting inflammation to depression. Therapeutic blockade of IL-6 signaling has proven clinically effective in treating many diseases including RA, Castleman’s disease, juvenile idiopathic arthritis, and giant cell arteritis, among others, highlighting the importance of tightly regulated IL-6 levels in immune homeostasis^21^. While extensive work has focused on IL-6 signaling and downstream effects across diverse cell types, far less is known about the upstream mechanisms governing IL-6 production, particularly in response to endogenous neuromodulators such as dopamine.

Emerging data link metabolic state and inflammatory output and highlight mitochondrial function as a central regulator of innate immune signaling^22,23^. Mitochondrial stress, altered dynamics, and mitochondrial DNA (mtDNA) release have all been implicated in the activation of innate immune pathways and production of inflammatory cytokines, including IL-6^24,25^. Indeed, mitochondrial integrity and altered metabolism are key regulators of macrophage inflammatory activation, where mitochondrial dysfunction can promote pro-inflammatory responses, such as cytokine production, which further amplify macrophage inflammatory signaling^26,27^. Prior data in human macrophages show that physiologic dopamine concentrations, including those induced by addictive drugs and some therapeutics, can activate several inflammatory pathways, including IL-6 release^17^. However, the mechanism(s) mediating the effects of dopamine on IL-6 remain unclear, and there are currently no data connecting the immunomodulatory effects of dopamine to the mitochondrial regulation of innate inflammation.

This study addresses this gap in our understanding of the role of dopamine in the regulation of innate immunity, defining the molecular mechanisms by which this catecholamine regulates IL-6. These data define a novel pathway by which dopaminergic signaling intersects with mitochondrial homeostasis and innate immune regulation in primary human monocyte-derived macrophages (hMDMs). Mechanistically, this response begins with dopamine-induced alterations in mitochondrial function, including mitochondrial membrane potential, excessive mitochondrial fission and a sustained rewiring of mitochondrial metabolism. The changes in fission are mediated by increased phosphorylation of dynamin-related protein 1 (DRP1), a GTPase that regulates mitochondrial fission. These events are mediated specifically by dopamine and promote the release of mtDNA into the cytoplasm. Cytosolic mtDNA is a potent endogenous danger signal, and it acts as a ligand for cyclic GMP-AMP synthase (cGAS), triggering STING activation and subsequent induction of inflammatory transcriptional programs, including NFκB-dependent production of IL-6. Pharmacologic blockade of dopamine-induced DRP1 activity and/or mtDNA release attenuated cGAS-STING activation and significantly reduced IL-6 secretion. Together, these findings define a mechanistic pathway linking dopamine signaling to IL-6-mediated inflammation, with potential implications for inflammatory states in which dopaminergic tone is altered.

## Methods

### Reagents

DMEM (high glucose) with GlutaMAX™ Supplement and pyruvate (#10569044, Gibco), fetal bovine serum (FBS; #MT35010CV, Gibco), human AB serum (#100-512-100, Gemini Bio), HEPES (BP2991), penicillin/streptomycin (#15140163, Gibco), and sterile diH₂O (#10977023, Invitrogen) were purchased from Thermo Fisher Scientific. Macrophage colony-stimulating factor (M-CSF; #315-02) was obtained from PeproTech. Dopamine hydrochloride (DA; #H8502, Sigma-Aldrich), lipopolysaccharide from *Escherichia coli* O55:B5 (LPS; #L2880, MilliporeSigma), carbonyl cyanide 4-(trifluoromethoxy)phenylhydrazone (FCCP; #0453, Tocris), 2′3′-cGAMP (#HY-12512, MedChemExpress), Mdivi-1 (#HY-15886, MedChemExpress), G140 (#HY-133916, MedChemExpress), H-151 (#HY-112693, MedChemExpress), ethidium bromide (EtBr; #HY-D0021, MedChemExpress), BAY 11-7082 (#S2913, Selleck Chemicals) were purchased from the indicated vendors.

Reagents were resuspended in sterile diH₂O or DMSO according to the manufacturers’ protocols, aliquoted as concentrated stock solutions, and stored at −20 °C or −80 °C protected from light. Fresh aliquots of all reagents were prepared from lyophilized powders every six months for all reagents, except dopamine hydrochloride, for which fresh aliquots were made every two months. Due to light sensitivity of some reagents, particularly dopamine hydrochloride, all reagent preparation and treatments were performed in reduced light conditions. For all experiments, working dilutions were freshly prepared in complete culture medium immediately prior to use. Where pharmacological inhibitors were employed, cells were pretreated for 60 min before stimulation. For DMSO-based reagents, the final DMSO concentration was kept constant across all conditions, and both vehicle-only and appropriate positive controls were included in every experiment.

### Human monocyte-derived macrophage (MDM) isolation and culture

Fresh human blood from de-identified healthy donors was obtained from the New York Blood Center (NYBC, Rye, NY, USA). Blood was obtained in accordance with the protocols approved by the Institutional Review Board (IRB) at the NYBC. All experiments with material isolated from human blood were performed using protocols approved by the Institutional Biosafety Committee at Drexel University (protocol No: 20984). Peripheral blood mononuclear cells (PBMCs) are isolated using Ficoll density gradient centrifugation techniques, as described previously^28^. Isolated PBMC were then centrifuged and resuspended in complete macrophage media consisting of DMEM-GlutaMAX supplemented with 10% heat-inactivated fetal bovine serum (FBS), 5% human AB serum, 1% HEPES, and 1% penicillin/streptomycin. Media was supplemented with macrophage colony-stimulating factor (M-CSF; 10 ng/mL).

To estimate the percentage of monocytes in the isolated PBMC, percentage of monocytes was determined by isolating monocytes from 10 million PBMCs using the Pan Monocyte Isolation Kit (#130-096-537, Miltenyi Biotec). PBMCs were resuspended and plated based on the estimated monocyte percentage, plating 100,000 monocytes per cm^2^ in each type of dish used (10 cm dishes; 6-, 12-, 48-, or 96-well plates). Following plating, monocytes were isolated via adherence selection and allowed to differentiate into macrophages over 6-7 days *in vitro*, yielding monocyte-derived macrophages (MDMs). At three days post-isolation and plating, cells were washed, and fresh complete macrophage media containing M-CSF was added, indicating enrichment of monocytes via adherence selection. Upon full maturation at 6 - 7 days *in vitro*, cells were used in experiments as described. Treatment durations and collection processes varied depending on the experimental design and are specified in subsequent Methods or Results sections.

### Cytokine and chemokine secretion analysis

Cytokine and chemokine secretion was assessed using culture supernatants collected from 48-well plates that were plated with 9.5 × 10^4^ monocytes per well. Supernatants were collected 24 hr after primary stimulation and IL-6 secretion was quantified using the AlphaLISA Human IL-6 Detection Kit (#AL223C, Revvity) as we have done previously ^17,28^. AlphaLISAs are high-throughput, bead-based assays; in these assays the IL-6 AlphaLISA had limit of detection of 10 pg/mL. A larger group of cytokines and chemokines were assessed using the MSD V-Plex Human Neuroinflammation Panel (#K15210D-1; Meso Scale Diagnostics), which we have shown to correlate with AlphaLISA analysis in these cells^28^. The MSD panel was used to measure different chemokines and cytokines listed in **Table. 1**. Plates were read on a MESO QuickPlex SQ 120 system and analyzed using MSD Discovery Workbench Software version 4.0 (Meso Scale Diagnostics). Donors with analyte levels below the limit of detection from Alphalisa were assigned a value of 10 pg/mL for fold change calculation and are indicated as not detectable (ND) in the corresponding figures.

### Western Blot

Protein expression and phosphorylation were assessed by Western blot analysis. Cells cultured in 6- or 12-well plates (9.5 x 10^5^ or 3.8 x 10^5^ monocytes per well, respectively) were lysed directly in wells using M-PER™ Mammalian Protein Extraction Reagent (Thermo Fisher Scientific) supplemented with 1% Halt™ Protease and Phosphatase Inhibitor Cocktail (Thermo Fisher Scientific). Lysates were collected by scraping and briefly sonicated to reduce viscosity. Protein concentration was determined using the Pierce™ BCA Protein Assay Kit (Thermo Fisher Scientific), and all samples were diluted to a final concentration of 1 µg/µL prior to downstream analysis. Lysates were stored at −80 °C until further use. Prior to immunoblotting, samples were diluted in 4X LI-COR loading buffer (#928-40004, LI-COR Biosciences), boiled for 10 min at 95 °C, and resolved on 4-12% Bolt™ Bis-Tris Plus gradient gels (Thermo Fisher Scientific) using MOPS running buffer. Proteins were transferred onto methanol-activated PVDF membranes (#88518, Thermo Fisher Scientific) using a wet-transfer system. Following transfer, membranes were stained with Revert™ Total Protein Stain (#926-11010, LI-COR Biosciences) and imaged on a LI-COR Odyssey imaging system for total protein-based normalization. Membranes were subsequently destained according to the manufacturer’s instructions and blocked in 3 - 5% bovine serum albumin (BSA; #BP160010, Fisher Scientific) in TBS-T for 1 hr at room temperature (RT) on a rotating shaker.

Following blocking, membranes were then incubated overnight at 4 °C with primary antibodies diluted in 3-5% BSA/TBS-T. Primary antibodies used included phospho-DRP1 (Ser616; #3455, Cell Signaling Technology, RRID: AB_2085352), total DRP1 (#8570, Cell Signaling Technology, RRID: AB_10950498), cGAS (D1D3G; #15102, Cell Signaling Technology, RRID: AB_2732795), phospho-STING (Ser366; E9A9K; #50907, Cell Signaling Technology, RRID: AB_2827656), total STING (D2P2F; #13647, Cell Signaling Technology, RRID: AB_2827656), phospho-TBK1 (Ser172; #PA5-105919, Thermo Fisher Scientific, RRID: AB_2817318), total TBK1 (#703154, Thermo Fisher Scientific, RRID: AB_2848223), phospho-IRF3 (Ser386; #MA5-35850, Thermo Fisher Scientific, RRID: AB_2849749), total IRF3 (#703682, Thermo Fisher Scientific, RRID: AB_2784599), phospho-AKT (Ser473; D9E; #4060, Cell Signaling Technology, RRID: AB_2315049), total AKT (#9272, Cell Signaling Technology, RRID: AB_329827), phospho-p38 MAPK (Thr180/Tyr182; D3F9; #4511, Cell Signaling Technology, RRID: AB_2139682), total p38 MAPK (#9212, Cell Signaling Technology, RRID: AB_330713), phospho-p44/42 MAPK (Erk1/2; Thr202/Tyr204; D13.14.4E; #4370, Cell Signaling Technology, RRID: AB_2315112), total p44/42 MAPK (Erk1/2; L34F12; #4696, Cell Signaling Technology, RRID: AB_390780), phospho-GSK3β (Ser9; 5B3; #9323, Cell Signaling Technology, RRID: AB_2115201), total GSK3β (27C10; #9315, Cell Signaling Technology, RRID: AB_490890), Parkin (#4211, Cell Signaling Technology, RRID: AB_2159920), PINK1 (#6946, Cell Signaling Technology, RRID: AB_11179069), BNIP3/Nix (#12396, Cell Signaling Technology, RRID: AB_2688036) and LC3B (#3868, Cell Signaling Technology, RRID: AB_2137707). The following day, membranes were washed three times with TBS-T and incubated with appropriate HRP-conjugated secondary antibodies, diluted in 5% milk, for 1 hr at RT. After additional washes (three times with TBS-T followed by a final wash with TBS), signals were detected using SuperSignal™ West Pico PLUS (#34580, Thermo Fisher Scientific) or SuperSignal™ West Femto Maximum Sensitivity Substrate (#34096, Thermo Fisher Scientific), as appropriate. For sequential detection of phosphorylated and total forms of the same protein, membranes were stripped by 0.25% glycine, 2% SDS, and 0.2% HCl (v/v) diluted in miliQ-H₂O for 8 min at RT and were washed thoroughly with TBS, and re-blocked in 3-5% BSA for 1 hr at RT before incubation with the next primary antibody overnight at 4 °C. This procedure was repeated as needed, with membranes not subjected to more than three stripping cycles. Phosphorylated and total protein expression levels were quantified using Image Studio™ Lite software (LI-COR Biosciences) and normalized to total protein expression or total protein stain levels, as appropriate.

### Spot Blot

Phosphorylation of the JAK/STAT pathway was assessed using the RayBiotech RayBio C-Series Human JAK/STAT Pathway Phosphorylation Array C1 (#AAH-JAKSTAT-1-8). These experiments were completed via the manufacturer’s protocol. Briefly, cells cultured in 10cm dishes (2.5 x 10^6^ monocytes per dish), treated with vehicle or dopamine for 30 min, then washed once with 1X PBS and then lysed directly in the dishes using 250µL of the provided Cell Lysis Buffer supplemented with the provided protease and phosphatase inhibitors. The cells in lysis buffer were pipetted up and down then rocked gently in plates at 4°C for 30 min. The plates were scraped and then the cells were collected, transferred to microcentrifuge tubes and centrifuged at 4°C at 14,000 x g for 10min. The lysates were then aliquoted in 50µL aliquots and stored at -80°C until use. Protein concentration was determined using the Pierce™ BCA Protein Assay Kit (Thermo Fisher Scientific), and 100µg of total protein was used for the assay. The assay was completed as recommended by the manufacturer with sample incubation at 5 hr at RT, detection antibody cocktail incubation overnight at 4°C , and the HRP-Anti-Rabbit IgG incubation for 2 hr at RT. All steps were performed using a rotating/rocking shaker at approximately 1 cycle/sec. Spot blots were imaged using the BioRad QuantityOne imaging system with a 2 sec exposure time. Densitometry of the spot blot was quantified using Image Studio™ Lite software (LI-COR Biosciences) and then analysis was completed by using the RayBiotech Excel-based Analysis Sheet.

### Global Proteomics and Phospho-site analysis

Lysates were collected as described for Western blotting. For each condition, five replicates were run 0.5 cm into a preparative SDS-gel and subjected to in-gel trypsin digestion. Tryptic digests were resuspended in 0.015% dodecyl maltoside, 0.1% trifluoroacetic acid before analysis. LC-MS/MS analysis was performed using an Orbitrap Astral mass spectrometer (Thermo Fisher Scientific) coupled to a Vanquish Neo UPLC system (Thermo Fisher Scientific). Chromatographic separation was carried out at 45 °C with a flow rate of 300 nL/min. Buffer A consisted of 0.1% formic acid in Milli-Q water, and buffer B consisted of 0.1% formic acid in acetonitrile. Peptides were loaded onto a trap column (100 Å, 75 μm i.d. x 2 cm packed with 3 μm C18 resin; Thermo Fisher Scientific), which was connected in line with a nanoEase M/Z Peptide BEH C18 nanocapillary analytical column (130 Å, 75 μm i.d. x 25 cm, 1.7 μm particle size; Waters). Separation was achieved using a 40 min linear gradient: 5-30% B over 25 min, 30-40% B over 6 min, 40-80% B over 2 min, followed by a 7 min hold at 80% B before re-equilibration.

MS data were acquired in data-independent acquisition mode. An MS1 scan was collected every 0.6 s in the Orbitrap at 240,000 resolution. Ions were injected for 3 ms or until an AGC target of 5e6 was reached. Precursor ions within a mass range of 380-980 m/z were collected in 2 m/z isolation windows with a maximum injection time of 3 ms or until an AGC target of 5e4 ions was reached. Precursors were fragmented using 25% normalized collisional energy. MS2 scans from 110 - 2000 m/z were collected in the Astral mass analyzer.

For data analysis, MS RAW files were searched with DIA-NN (v2.2.0)^29^ against a spectral library generated from the SwissProt human database with curated isoforms (UP000005640, downloaded 4-25-2025). N-terminal acetylation, N-terminal methionine excision, and methionine oxidation were set as variable modifications. Cysteine carbamidomethylation was set as a static modification. Data were searched with full tryptic specificity, allowing for 2 missed cleavages and 1 variable modification, a peptide length of 7-50 amino acids, a precursor charge of 1-4, and a scan window of 15. MS1 and MS2 mass accuracy were set to 5.0 and 10.0 ppm, respectively. The match-between-runs feature was enabled, and the cross-run normalization feature was disabled.

Contaminants, proteins identified by a single peptide, and proteins with less than 4 valid values in any condition were removed from the filtered dataset. Protein abundance was normalized using the quantile method, implemented in NormalyzerDE^30^. Differential expression analysis was performed using LFQ Analyst^31^. For proteins with no valid values, missing values were imputed with the minimum value of the dataset. For proteins with one or two valid values, missing values were imputed with the minimum peptide abundance. The FDR level was controlled using the Benjamini-Hochberg correction. Proteins with a fold change > 2 and an adjusted p-value < 0.05 were considered significant.

For the analysis of phosphosites, the search was repeated with the following modifications to the settings: phosphorylation of serine, threonine, and tyrosine residues was added as a variable modification, and a maximum of two variable modifications were allowed. The phosphosite matrix with a localization confidence of greater than 90% was used for further analysis. Contaminants and phosphosites with less than four valid values in any condition were removed from the filtered dataset.

### High Content Imaging and Analysis (Nuclear Translocation)

Nuclear translocation of phosphorylated NF-κB (p-NF-κB) was assessed using our previously established high-content imaging protocols^28^. Briefly, Nunc™ Microwell 96-well optical-bottom plates (Thermo Fisher Scientific) were plated with 1.5 x 10^4^ monocytes per well and were treated with dopamine or vehicle control, or LPS (10ng/ml), for 1 hr. Cells were then fixed with 4% paraformaldehyde (PFA) for 10-15 min at RT and washed with PBS. At this stage, plates were either processed immediately or stored at 4°C for up to one week prior to immunofluorescence staining. Cells were permeabilized with 0.1% Triton™ X-100 in PBS for 5 min, washed with PBS, and blocked for 1 hr at RT using blocking buffer containing 1% bovine serum albumin (BSA) and 300 mM glycine in PBS with 0.1% Tween-20. Following blocking, cells were incubated overnight at 4 °C with primary antibodies against phospho-NF-κB p65 (Ser536; #8242, Cell Signaling Technology, RRID: AB_10859369), diluted in blocking buffer. Following primary incubation, cells were washed with PBS and incubated for 1 hr at RT with Alexa Fluor 488 goat anti-rabbit 2° antibody (#A-11008, Thermo Fisher Scientific, RRID: AB_143165) diluted in blocking buffer. Nuclei were counterstained with DAPI, and the plasma membrane was labeled using CellMask™ Deep Red (ThermoFisher Scientific).

Images were acquired using a CellInsight™ CX7 High-Content Screening Platform (Thermo Fisher Scientific) with a 20X objective. Each experimental condition was performed in triplicate wells to minimize technical variability. For each well, 1,000 - 1,500 cells were imaged and analyzed using defined parameters (**Supp Table. 1**). Image analysis was performed using HCS Studio software, where binary masks were generated to define nuclear and cytoplasmic compartments. The intensity of p-NF-κB signal within each compartment was quantified, and nuclear translocation was calculated as the ratio of nuclear to cytoplasmic intensity for each cell.

### Confocal Imaging and Analysis of Mitochondrial Membrane Potential

To assess mitochondrial membrane potential, cells were stained with JC-1 dye (#T3168, Thermo Fisher Scientific). Lyophilized JC-1 was initially dissolved in DMSO to a stock concentration of 25 mg/mL, aliquoted, and stored at −20°C protected from light. On the day of imaging, freshly thawed JC-1 was diluted in phenol red-free DMEM to a final concentration of 10 µg/mL and incubated with cells for 10 min at 37 °C. Cells were then washed twice with phenol red-free DMEM and imaged live using an Olympus FV3000 confocal microscope with a 20X objective (Numerical Aperture = 0.75), equipped with temperature-, humidity-, and CO₂-controlled environmental chamber. Images were acquired from three wells per condition, with three distinct regions of interest (ROIs) imaged per well. Image analysis was performed using FIJI software by quantifying fluorescence intensity in the 488 nm and 555 nm channels, corresponding to monomeric JC-1 (low mitochondrial membrane potential) and J-aggregate JC-1 (high mitochondrial membrane potential), respectively.

### Confocal Imaging and Analysis of Mitochondrial Morphology

To evaluate changes in mitochondrial morphology analysis, lyophilized MitoTracker™ Orange CMTMRos (#M7510, Thermo Fisher Scientific) was initially dissolved in DMSO to generate a 1 mM stock solution, aliquoted, and stored at −20 °C protected from light. On the day of the experiment, the dye was diluted in pre-warmed, phenol red-free DMEM to a final concentration of 250 nM, and live cells were incubated with the dye for 20 min at 37 °C in a humidified incubator. Cells were subsequently fixed with 4% paraformaldehyde (PFA) for 15 min at RT and washed with PBS. Nuclei were counterstained with DAPI. Three-dimensional confocal imaging of mitochondria was performed using an Olympus FV3000 confocal microscope with a 30X oil-immersion (Numerical Aperture = 1.05) Z-stack images were acquired using a step size of 0.15 µm. For each treatment condition, images were collected from three independent wells, with 25 randomly selected fields imaged per condition. Image analysis was performed in a blinded manner using the MitoAnalyzer plug-in in FIJI (ImageJ) software to quantify mitochondrial morphology parameters^32^.

### Confocal Imaging and Analysis of Mitophagy

To assess mitophagy, fixation and staining procedures were performed as described above. Cells were incubated with primary antibodies against LAMP1 (lysosomal marker, #9091, Cell Signaling Technology, RRID: AB_2687579) and TOMM20 (mitochondrial outer membrane marker, #ab289670, Abcam, RRID: AB_2943038), followed by incubation with appropriate Alexa Fluor-conjugated secondary antibodies (Alexa Fluor 488 goat anti-rabbit and corresponding anti-rat secondary antibodies; Thermo Fisher Scientific, RRIDs: AB_143165 and AB_141778). Nuclei were counterstained with DAPI. 3D confocal imaging of mitochondria was performed using an Olympus FV3000 confocal microscope with a 30X oil-immersion (Numerical Aperture = 1.05), and Z-stack images were acquired using a step size of 0.15 µm. Image analysis was performed on Z-projection images using the Coloc2 plug-in in FIJI (ImageJ) software to quantify mitochondrial morphology parameters.

### Flow Cytometry

Mitochondrial superoxide production was assessed in live hMDMs using flow cytometry. Following treatment with dopamine or vehicle, cells were gently detached using TrypLE™ Express Enzyme (#12605028, Thermo Fisher Scientific), collected, and stained with MitoSOX™ Red mitochondrial superoxide indicator (#M36008, Thermo Fisher Scientific) for 30 min at 37 °C in the dark. To discriminate live and dead cells, Zombie NIR™ fixable viability dye (#423105 , BioLegend) was included. A dead-cell control was generated by heat-killing a subset of cells at 55 °C for 10 min and mixing them 1:1 with live cells. Additional controls included unstained samples and single-stained controls for accurate gating and compensation. Following staining, samples were acquired on a CytoFLEX™ flow cytometer (Beckman Coulter). Data were analyzed using FlowJo software, with mitochondrial superoxide levels quantified in the live cell population after exclusion of debris and dead cells.

### Analysis of Cellular Respiration

Analysis of changes in cellular respiration were performed using an Agilent Seahorse XFe96 Analyzer. Oxygen consumption rate (OCR; pmol O₂/min) and extracellular acidification rate (ECAR; mpH/min) were measured in live, treated hMDMs. Cells were seeded at a density of 1.5 x 10^4^ monocytes per well in Seahorse XF96 cell culture microplates (Agilent), a density previously optimized across multiple plating conditions. hMDMs were treated with dopamine, along with appropriate vehicle and positive controls (FCCP: #0453/10, Tocris), for durations specified in the Results section. Following treatment, cells were washed twice with Seahorse XF DMEM assay medium supplemented with glucose, pyruvate, and L-glutamine, and then incubated for 60 min in a non-CO₂ incubator at 37 °C to allow temperature and pH equilibration. The Seahorse XF Real-Time ATP Rate Assay Kit (#103592-100, Agilent) and the Seahorse XF Cell Mito Stress Test Kit (#103015-100, Agilent) were used according to the manufacturer’s instructions. For the ATP rate assay, injection ports were loaded with oligomycin (first injection, 1.5 µM) followed by a combination of rotenone and antimycin A (second injection, 0.5 µM). For the Mito Stress Test assay, FCCP (1 µM) was added as a second injection, while rotenone and antimycin A were added as a third injection. The sensor cartridge was hydrated, calibrated, and equilibrated in parallel using pre-warmed calibration buffer. Following completion of the assay, cells were stained with Hoechst nuclear dye (Thermo Fisher Scientific) and imaged using an ImageXpress Pico Automated Cell Imaging System (Molecular Devices). Cell counts were determined per well and used to normalize OCR and ECAR measurements to cell number.

### Subcellular Fractionation

To interrogate changes specific to proteins in the cytosol and mitochondria, subcellular fractionation was used to isolate cytosolic and mitochondrial fractions from hMDMs plated in 10 cm dishes at 3 x 10^6^ cells per dish. Following experimental treatments, cells were gently detached using TrypLE™ Express Enzyme (#12605028, Thermo Fisher Scientific), collected, and subjected to subcellular fractionation using the Mitochondria/Cytosol Fractionation Kit (#ab65320, Abcam) according to the manufacturer’s protocol. The cytosolic fraction was collected using the kit-provided cytosolic extraction buffer and further diluted with appropriate lysis buffers depending on downstream applications. For DNA isolation, cytosolic fractions were diluted at a 2:1 ratio with DNA/RNA Lysis Buffer (#D7001-1, Zymo Research). For immunoblotting, cytosolic fractions were mixed with 2X Laemmli sample buffer (#1610737, Bio-Rad). The mitochondrial fraction was collected as a pellet and resuspended in the appropriate lysis buffer depending on the experimental application, as described above. Protein concentrations were determined using the BCA assay prior to downstream analyses. To validate the purity of subcellular fractions, samples were analyzed by Western blotting following the protocol described above. Cytosolic fractions were probed for β-actin (13E5; #4970S, Cell Signaling Technology, RRID: AB_2223172) as a cytosolic marker, while mitochondrial fractions were validated by probing for cytochrome c oxidase subunit IV (COX IV; 4D11-B3-E8; #11967, Cell Signaling Technology, RRID: AB_2797784).

### Quantitative PCR

Total RNA was isolated from cells using the Quick-DNA/RNA™ Miniprep Kit (Zymo Research) according to the manufacturer’s instructions. RNA concentration and purity were assessed using a NanoDrop One spectrophotometer (Thermo Fisher Scientific). For each donor, 1µg of total RNA was reverse-transcribed into cDNA using the High-Capacity cDNA Reverse Transcription Kit (#4368814, Applied Biosystems). Quantitative PCR was performed using 12.5ng of cDNA per reaction, with each target gene analyzed in triplicate technical replicates. mRNA expression levels of dopamine receptor genes (DRD1-DRD5; assay IDs: Hs00265245_s1, Hs00241436_m1, Hs00364455_m1, Hs00609526_m1, Hs00361234_s1), as well as IFIT1 (Hs01675197_m1) and ISG15 (Hs01921425_s1), were quantified using TaqMan™ Fast Universal PCR Master Mix and gene-specific TaqMan™ assays (Thermo Fisher Scientific). Expression of 18S (#4319413E, Thermo Fisher Scientific) used as the endogenous housekeeping control and only data from samples with 18s values between 8-13 were included. All TaqMan reagents were obtained from Thermo Fisher Scientific. To assess the release or leakage of mitochondrial DNA (mtDNA) into the cytoplasm, genomic DNA (gDNA) was extracted from the cytoplasmic fraction using the Quick-DNA/RNA™ Miniprep Kit (Zymo Research). DNA concentration was measured using a NanoDrop One spectrophotometer. Quantitative PCR was performed using 0.5ng of gDNA per reaction, with each gene run in triplicate technical replicates, using SYBR™ Green Master Mix (Bio-Rad) and primers targeting Cytochrome c oxidase (F: 5’GCCCCAGATATAGCATTCCC-3’ and R: 5’-GTTCATCCTGTTCCTGCTCC-3’). Nuclear 18S (F: 5’-TAGAGGGACAAGTGGCGTTC-3’ and R: 5’CGCTGAGCCAGTCAGTGT-3’) rRNA was used as the reference gene. Relative gene expression levels were calculated using the comparative Ct (ΔΔCt) method, with values first normalized to 18S expression levels and then expressed relative to the control condition.

### Principal Component Analysis (PCA)

The six-feature panel (DRD1-DRD5 and IL-6 fold change) was used for dimensionality reduction. Only donors with complete data for all six features were retained. DRD1-DRD5 were log₁₀-transformed after adding a small constant equal to half the minimum positive value. The IL-6 fold change was log₂-transformed with the same pseudo-count rule (appropriate for a ratio). The transformed features were then standardized to zero mean and unit variance using sklearn.preprocessing.StandardScaler (scikit-learn=1.4.0). PCA was performed on the standardized matrix using sklearn.decomposition.PCA with default settings (all components retained). Unsupervised hierarchical clustering was performed on the same standardized matrix (log₁₀ DRD1-DRD5; log₂ fold change). Pairwise Euclidean Distances were computed using scipy.spatial.distance.pdist (scipy=1.11.0) and Ward linkage was computed with scipy.cluster.hierarchy.linkage (method ’ward’). The optimal number of clusters K was selected by evaluating silhouette scores (sklearn.metrics.silhouette_score, metric ’euclidean’) for K ∈ {2, 3, 4, 5, 6} using sklearn.cluster.AgglomerativeClustering with Ward linkage. The value of K that maximized the silhouette score was retained. Final cluster labels were assigned with the chosen K and merged onto the full donor table by Donor identifier.

### Statistics

All data were assessed for normality by assessing skewness and testing for normal or lognormal distribution to inform the selection of appropriate statistical analyses. Data were primarily normalized to the mean of the vehicle control to account for inter-donor variability inherent to primary human cell experiments and to report treatment-induced fold changes. Technical outliers were identified using the ROUT method (Q = 0.1%) and excluded where applicable. *Post hoc* comparisons were performed as appropriate. All statistical analyses were conducted using GraphPad Prism version 10.2 (GraphPad Software, La Jolla, CA), with statistical significance defined as p < 0.05.

## Results

### Dopamine induces IL-6 secretion in human monocyte-derived macrophages in a receptor-dependent manner

Our prior work demonstrated that dopamine increases the production and secretion of several cytokines, including IL-6 and IL-1β, in human myeloid cells^9,10,17^. To define the mechanistic basis by which dopamine induces IL-6 secretion, hMDM were differentiated from isolated human PBMC and treated with dopamine, as well as lipopolysaccharide (LPS) and vehicle (H_2_O) as positive and negative controls (**Fig. 1A)**. Based on prior dose-response studies, hMDMs were treated with 1 µM dopamine, a concentration that produces maximal IL-6 secretion as well as nuclear translocation of NF-κB^10,17^. Culture supernatants were collected 24 hr after stimulation, and IL-6 levels were quantified using AlphaLISA. Dopamine treatment resulted in a significant, 2.1-fold increase in IL-6 secretion relative to vehicle (**Fig. 1B, Supp. Fig. 1A**). Dopamine-induced increases in IL-6 were confirmed by multiplex MSD V-Plex assays. These assays highlight the donor-to-donor variability in primary human samples, while also showing increases in IL-7 and increasing trends in the secretion of IL-12p70, IL-2, IL-4, IL-17 and IL-13. No significant changes were observed in the secretion of the other cytokines tested, indicating selectivity in dopamine-driven cytokine responses (**Table. 1**). These effects were specific to dopamine, as treatment of a subset of donors with dopamine metabolites such as 3,4-dihydroxyphenylacetic acid (DOPAC) and homovanillic acid (HVA), as well as levodopa (L-DOPA), an essential dopamine precursor did not induce IL-6 secretion. Moreover, other monoamine neurotransmitters, including serotonin (SER) and norepinephrine (NE) also failed to elicit similar responses, further supporting a dopamine-specific signaling mechanism (**Fig. 1C)**.

**Figure 1.**
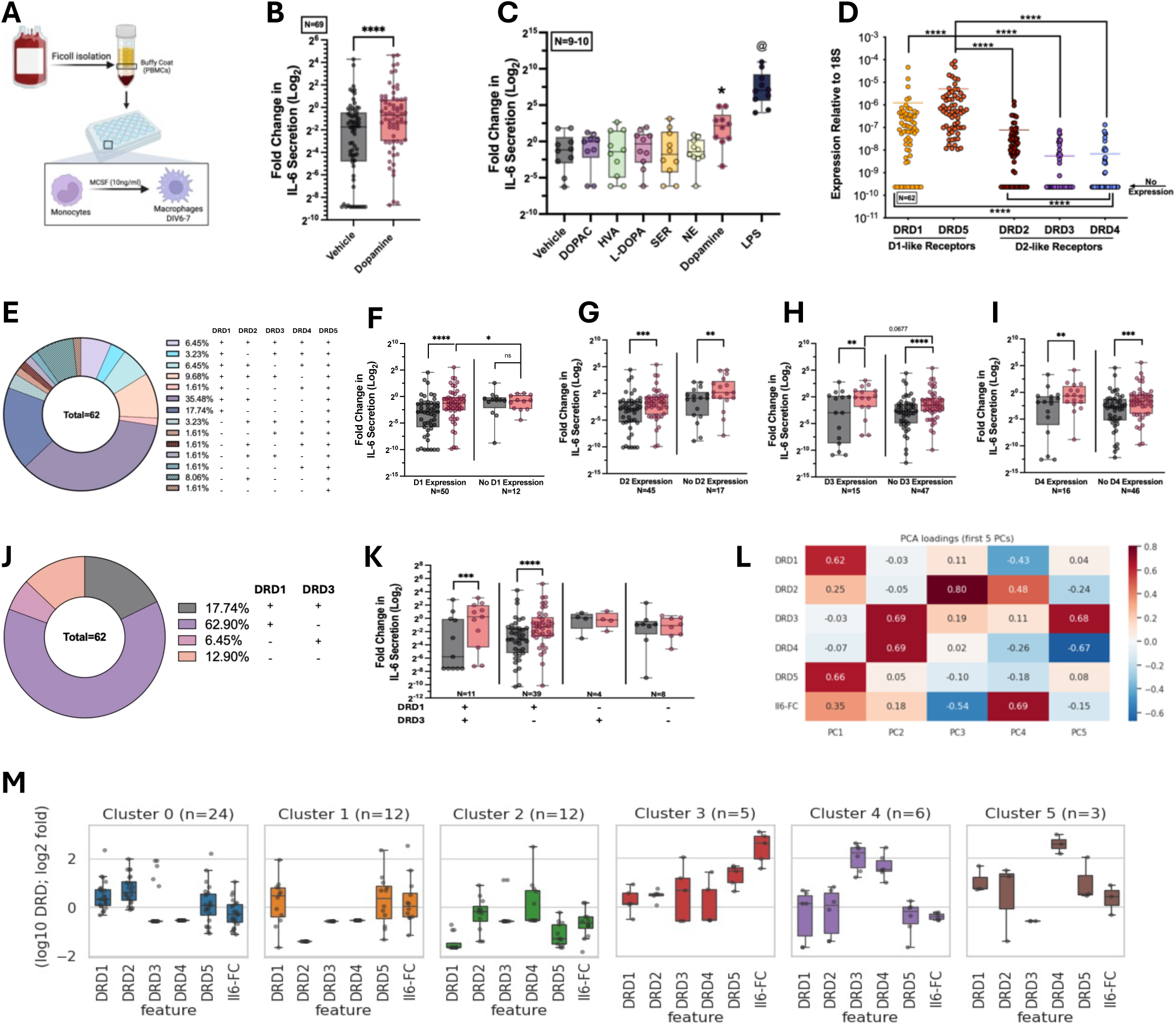
Dopamine induces IL-6 secretion in human monocyte derived macrophages in a receptor-dependent manner. (A) PBMCs were isolated from fresh human blood and plated according to monocyte counts. Monocytes were further enriched by adherence-based isolation and differentiated into hMDMs over 6-7 days in the presence of M-CSF. Fully differentiated hMDMs were then treated with dopamine or appropriate controls for downstream experiments. Created on Biorender.com. (B) Dopamine (1 μM) treatment induces a significant increase in IL-6 secretion 24 hr post-treatment, as measured by AlphaLISA (Biological replicates: N=69). Data are presented as box and whisker plot (min to max). See Supplemental Figure 1A. (C) This effect is specific to dopamine (1 μM), as hMDMs were also treated with dopamine metabolites (DOPAC, 1 μM; HVA, 1 μM), the dopamine precursor (L-DOPA, 1 μM), and other catecholamines (SER, 1 μM; NE, 1 μM). LPS (10 ng/mL) was included as a positive control, and IL-6 secretion was measured by AlphaLISA 24hr post-treatment. (Biological replicates: N=9-10). Data are presented as box and whisker plot (min to max). (D) Untreated hMDMs express multiple dopamine receptors, as measured by qPCR, including DRD1 and DRD5 (D1-like receptors) as well as DRD2, DRD3, and DRD4 (D2-like receptors), (Biological replicates: N=62). Individual values are shown with line demonstrating mean. (E) DRD expression patterns vary in different donors, shown in pie chart (Biological replicates: N=62). (F-I) Donors from Fig 1A were stratified based on the presence or absence of individual DRDs (Fig 1D), and dopamine-induced IL-6 secretion levels were compared between groups to assess potential correlations between each individual DRD expression and dopamine-induced IL-6 production. (J) Donors were stratified based on the presence or absence of DRD1 and DRD3 expression, and the proportions of each population are shown in a pie chart. (K) The fold change in IL-6 secretion in response to dopamine exposure (measured by AlphaLISA) was compared across the populations defined in Fig 1.J. (L) Heatmap of PCA loadings for the first five principal components. Each row is one panel feature (DRD expressions and fold change in IL6 secretion in response to dopamine); color encodes the signed loading magnitude (red = positive, blue = negative), (Biological replicates: N=62). See Supplemental Figure 1B and C. (M) Unsupervised clustering was performed based on PC1 and PC2. Each subplot corresponds to one of the six clusters (cluster ID and n shown in subtitle). Boxes show the median and IQR; individual donor values are overlaid as jittered points. Features with large z-score deviations relative to the other clusters indicate the primary drivers of cluster identity (Biological replicates: N=62). Paired comparisons were analyzed using a paired *t*-test (non-parametric: Wilcoxon test) in panel B,C and K. Comparisons between ±DRDs groups were performed using an unpaired *t*-test (Mann-Whitney test) in panel F-I. Significance levels are indicated as follows: *p* < 0.05 (* or @), *p* < 0.01 (**), *p* < 0.001 (***), and *p* < 0.0001 (****).

As dopamine primarily exerts its effects through dopamine receptor signaling, we quantified expression of dopamine receptor transcripts in hMDMs from 62 donors using quantitative PCR (qPCR). Consistent with prior findings^9^, all donors expressed DRD5, whereas expression of DRD1-4 varied between individuals (**Fig. 1D**). Based on these expression patterns, donors were classified into 14 distinct groups, with the most prevalent group (35.48%) characterized by expression of DRD1, DRD2, and DRD5, and absence of DRD3 and DRD4 (**Fig. 1E**). Stratification based on individual receptor expression revealed that donors expressing DRD1 exhibited significantly higher dopamine-induced IL-6 secretion (**Fig. 1F**). In contrast, donors with or without DRD2 expression showed comparable IL-6 induction following dopamine treatment (**Fig. 1G**). Donors expressing or lacking DRD3 both demonstrated robust IL-6 responses, with a trend toward increased responses in DRD3-expressing donors (**Fig. 1H**). Similar to DRD2, DRD4 expression did not influence responsiveness, as donors with or without DRD4 exhibited comparable IL-6 secretion (**Fig. 1I**). Given the differential responses associated with DRD1 and DRD3, donors were further stratified based on the presence or absence of these two receptors. The majority of donors (62.9%) expressed DRD1 but not DRD3, whereas 17.75% expressed both receptors. Donors lacking DRD1 but expressing DRD3 or lacking both receptors comprised 6.45% and 12.90% of the population, respectively (**Fig. 1J**). Comparison of dopamine-induced IL-6 secretion across these groups confirmed donor-dependent variability, with the highest responses observed in donors expressing DRD1. Expression of DRD3 alone did not correlate with a dopamine mediated increase in IL-6, suggesting that DRD1 expression is a primary determinant of responsiveness, whereas DRD3 plays a more modulatory role (**Fig. 1K**).

Principal component analysis (PCA) of DRD expression and IL-6 response to dopamine (fold change) revealed that Principal component 1 (PC1) is primarily driven by DRD1 and DRD5 expression, with additional contribution from IL6 response and DRD2, whereas PC2 is dominated by DRD3 and DRD4 expression, reflecting distinct receptor-specific expression patterns (**Fig. 1L**). Unsupervised clustering identified six clusters (Cluster 0-5), each exhibiting unique DRD expression profiles and differential responsiveness to dopamine (**Supp. Fig. 1B**). Cluster-specific analysis revealed distinct receptor patterns. Cluster 0 (N=24), the largest group, displayed moderate expression of DRD1, DRD2, and DRD5 with relatively low DRD3/DRD4 levels and modest IL6 response. Cluster 1 (N=12) showed a similar but more variable pattern, with slightly elevated DRD5 expression, low expression of D2-like receptors (DRD2/DRD3/DRD4) expression and heterogeneous IL6 responses. In contrast, Cluster 2 (N=12) exhibited higher DRD4 expression with reduced D1-like receptor levels (DRD1/DRD5) and diminished IL-6 response. Cluster 3 (N=5) demonstrated the strongest IL-6 induction alongside elevated DRD1, DRD3 and DRD5 expression. Cluster 4 (N=6) was characterized by high DRD3/DRD4 expression with lower DRD1/DRD5 and diminished IL6 response, whereas Cluster 5 (N=3) showed elevated DRD1 and DRD4 expression but relatively low IL6 induction (**Fig. 1M, Supp. Fig. 1C**). Demographic stratification of our donor set revealed that both male and female donors responded to dopamine stimulation, with comparable increases in IL-6 production. However, baseline IL-6 secretion in hMDMs from female donors was higher (**Supp Fig. 1D**). Similarly, macrophages from both under 50-year-old and over 50-year-old donors showed robust dopamine-induced IL-6 secretion, but donors over 50 years of age exhibited higher baseline IL-6 levels (**Supp Fig. 1E**). Bivariate analysis of donor age and dopamine-induced IL-6 response (fold change) demonstrated that dopamine consistently induced IL-6 secretion across all age and gender groups, independent of these demographic variables (**Supp Fig. 1F**).

### Dopamine Induces Time-Dependent Changes in Global Protein Abundance and Phosphosite-Specific Signaling in Human Macrophages

To define the molecular substrates underlying dopamine-induced increases in IL-6 secretion, we performed global proteomic analysis on hMDMs derived from five matched donors treated with vehicle or dopamine (1 μM) for 1 hr or 6 hrs. Analysis of these data show that 1 hr of dopamine treatment induced significant increases in the abundance of several proteins, including HLA-DRB4, TMBIM4, MGST3, and NABP2 (**Fig. 2A**). Increases in HLA-DRB4 suggest a heightened activation state^33^, while increased TMBIM4, an endoplasmic reticulum (ER)/Golgi-associated protein involved in calcium regulation^34^, and MGST3, linked to oxidative stress responses, suggest early alterations in cellular stress pathways, including those related to organelle homeostasis^35^. The induction of NABP2, a DNA-binding protein involved in DNA damage responses^36^, further supports engagement of cellular stress signaling networks at early time points.

**Figure 2.**
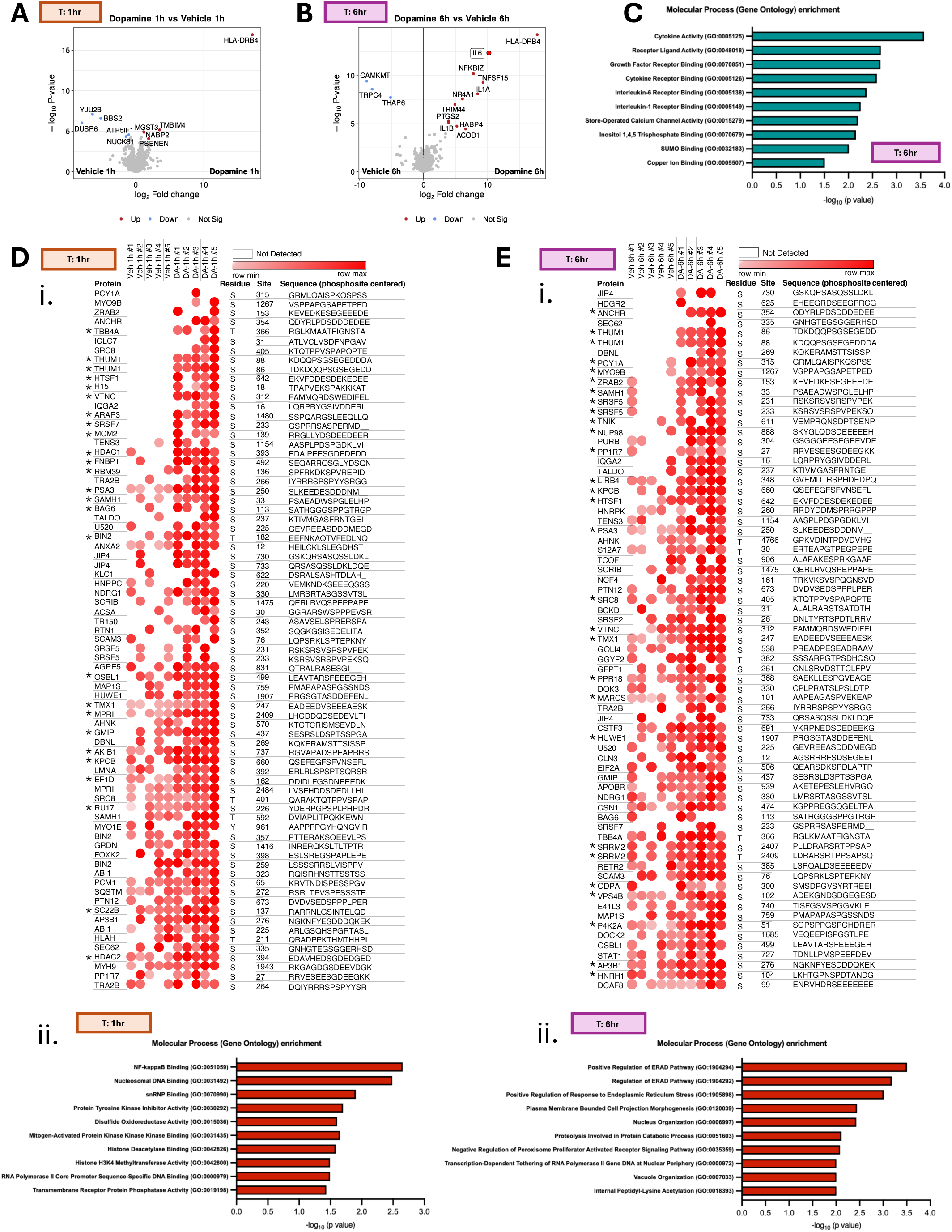
Dopamine Induces Time-Dependent Changes in Global Protein Abundance and Phosphosite-Specific Signaling in Human Macrophages. (A,B) hMDMs from five distinct donor groups (biological replicates: N=5) were treated with vehicle or dopamine (1 μM) and collected at 1 or 6 hr post-treatment. Samples were analyzed for global proteomics, and proteins with significantly altered abundances are shown in volcano plots. See Supplemental Figure 2A and B. (C) Gene Ontology (GO) molecular process enrichment analysis was performed on proteins with significantly increased abundance following 6 hrs of dopamine treatment. (D-E) Phospho-site analysis was performed for both vehicle vs dopamine (1 hr:D and 6 hr:E) groups. (biological replicates: N=5) (i) Heatmaps summarize proteins with ≥2-fold increased phosphorylation. * indicates proteins with statistically significant increases (p ≤ 0.05) within this group. See Supplemental Figure 2C-F. (ii) Proteins with significant changes were subjected to GO molecular process enrichment analysis, and enriched pathways are shown.

At later time points (6 hrs post-treatment), dopamine induced a broader and more robust proteomic response, with the most robust changes associated with IL-6 production and expression of HLA-DRB4. There was also increased abundance of inflammatory mediators such as NFKBIZ, TNFSF15, IL1A, IL1B, and NR4A1 (**Fig. 2B**). A small number of differences were observed between vehicle-treated groups across time points; however, these did not overlap with the experimental comparisons, indicating that the observed effects were attributable to dopamine (**Supp Fig. 2A**). Among the proteins with the largest increases in response to 6 hr dopamine treatment, NFKBIZ (IκBζ) is a known regulator of IL-6 transcription^37^, while IL1A/IL1B and TNFSF15 are consistent with activation of inflammatory signaling pathways^9,38^. Gene Ontology (GO) molecular process enrichment analysis of significantly upregulated proteins at 6 hr revealed enrichment in pathways associated with cytokine production and receptor-mediated inflammatory signaling, with specific activation of IL-6 mediated signaling pathways (**Fig. 2C**), corroborating the large increase in IL-6 seen between 1 hr of dopamine treatment and 6 hrs of dopamine treatment (**Supp Fig. 2B**).

As many signaling pathways are regulated by protein phosphorylation, we assessed phospho-site-specific changes by analyzing proteins exhibiting ≥2-fold increases in phosphorylation at 1 or 6 hr post-treatment. Significantly altered phospho-sites highlighted in heatmaps (**Fig. 2Di,Ei**). GO enrichment of these significantly regulated phosphoproteins demonstrated time-dependent pathway engagement, with NFκB-associated signaling pathways predominantly affected at 1 hr, and pathways related to the ER emerging at 6 hr (**Fig. 2Dii,Eii**). To further define functional connectivity, we performed STRING interaction enrichment analysis on proteins exhibiting ≥2-fold increases in phosphorylation, revealing interconnected protein networks (**Supp Fig. 2C,D**). Proteins were clustered based on the MCL method (**Supplementary Table. 2)**. Notably, 8 phospho-sites overlapped between the 1 hr and 6 hr time points, and these proteins were primarily associated with mitochondrial and ER compartments (**Supp Fig. 2E,F**), suggesting coordinated dysregulation of these organelles over time in response to dopamine exposure.

### Dopamine-induced IL-6 secretion is mediated by NF-κB and activation of the cGAS-STING pathway

Nuclear factor kappa B (NF-κB) is a central regulator of inflammatory gene expression, including IL-6. Our prior studies have shown that dopamine activates NF-κB, driving phosphorylation and degradation of IκB, leading to the release, phosphorylation, and nuclear translocation of active NF-κB^10^. Consistent with these observations, our global proteomics dataset revealed increased expression of NFKBIZ, suggesting engagement of NF-κB-dependent transcriptional programs. To confirm this, NF-κB activation was quantified by assessing the subcellular localization of phosphorylated NF-κB (p-NF-κB) in response to dopamine using immunofluorescence and high content imaging. As anticipated, dopamine induced a significant but modest (∼24%) increase in p-NF-κB nuclear translocation, at 1 hr post-treatment (**Supp Fig. 3A,B**). To determine whether dopamine-induced IL-6 secretion is mediated through this pathway, we pharmacologically inhibited NF-κB activation by pretreating hMDM with BAY 11-7082 (10 μM, BAY-11), an inhibitor of IκB phosphorylation^39^, for 1 hr prior to dopamine treatment. Inhibition of NF-κB signaling significantly reduced dopamine-induced IL-6 secretion compared with dopamine treatment alone (**Fig. 3A**), indicating that NF-κB activation is necessary for the dopamine-mediated increase in IL-6 production.

**Figure 3.**
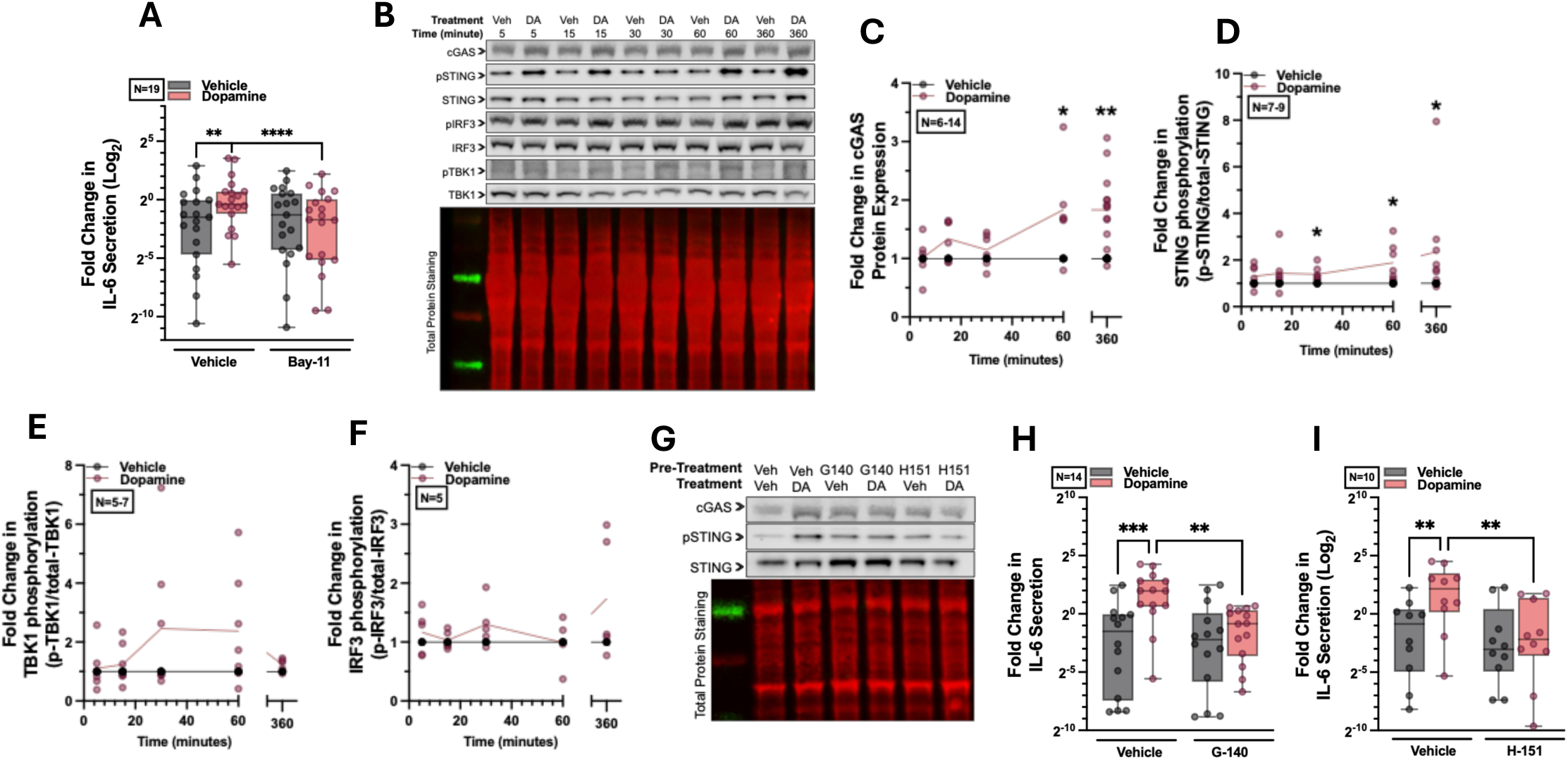
Dopamine-induced IL-6 secretion is mediated by NF-κB and activation of the cGAS-STING pathway. (A) Cells were pretreated (t = −1 hr) with BAY 11-7082 (10 μM) to inhibit NFκB activation. IL-6 secretion was measured 24 hrs after dopamine treatment using AlphaLISA, demonstrating that dopamine-induced IL-6 secretion is mediated through the NFκB pathway, (Biological replicates: N=19). Data are presented as box and whisker plot (min to max). See Supplemental Figure 3A and B. (B-F) Cells were treated with dopamine in a time-course experiment (5, 15, 30, 60, and 360 minutes) and assessed for cGAS protein expression (normalized to total protein expression, Biological replicate: N=5-14) as well as phosphorylation of STING (Biological replicate: N=7-9), TBK1 (Biological replicate: N=5-7), and IRF3 (Biological replicate: N=5), (normalized to the total expression level of each protein) by western blotting. Individual values are shown with line demonstrating mean. See Supplemental Figure 2J and K. (G-I) Cells were pretreated (t = −1 hr) with G-140 (1 μM) or H-151 (1 μM) to inhibit the cGAS-STING pathway, and the inhibitory effects were validated by western blotting. Cells were then treated with dopamine (t = 0). IL-6 secretion was measured 24 hrs post-treatment using AlphaLISA (Biological replicate: G-140:N=14 and H-151: N=10), demonstrating that activation of the cGAS-STING pathway contributes to dopamine-induced IL-6 secretion. Data are presented as box and whisker plot (min to max). Paired comparisons were analyzed using a paired *t*-test (non-parametric: Wilcoxon test) in panel C-F or paired one-way ANOVA (non-parametric: Friedman test) in panel A, H and I. Significance levels are indicated as follows: *p* < 0.05 (*), *p* < 0.01 (**), *p* < 0.001 (***).

To define the signaling events linking dopamine signaling to NF-κB, hMDMs were stimulated with dopamine (5 - 60 min) and assessed for activation of NF-κB associated pathways, including AKT, p38 MAPK, ERK1/2 (p42/44), GSK3β and STAT signaling, via immunoblotting^39,40^. Dopamine did not induce phosphorylation of these pathways within 1 hr of treatment (**Supp Fig. 3C-I**), suggesting that dopamine-mediated NF-κB activation occurs independently of these signaling cascades. The proteomic and phospho-proteomic analyses also showed enrichment of organelle-associated pathways, including those linked to the ER and mitochondrial compartments. Therefore, we assessed the potential involvement of intracellular stress signals involved in cytosolic danger sensing^41–43^. Emerging evidence links innate immune activation to cytosolic DNA sensing, prompting examination of the cyclic GMP-AMP synthase (cGAS)-stimulator of interferon genes (STING) pathway, a key signaling axis upstream of NF-κB activation in human myeloid cells^44,45^. Immunoblotting analysis of dopamine-treated hMDMs revealed a significant increase in cGAS protein levels 1 hr post-treatment, which persisted for at least 6 hrs (**Fig. 3B,C**). Consistent with this, dopamine treatment also induced STING phosphorylation (Ser366) at the same time points (**Fig. 3B,D**). Downstream of STING, dopamine induced an increasing trend in TBK1 phosphorylation in a subset of donors (**Fig. 3B,E**).

In addition to NF-κB, activation of the cGAS-STING pathway can promote phosphorylation and nuclear translocation of IRF3^46,47^, leading to transcription of type I interferon-stimulated genes. Despite robust activation of cGAS and STING, dopamine did not induce IRF3 phosphorylation (**Fig. 3B,F**). Corroborating the lack of IRF3 activity, dopamine did not upregulate IRF3-dependent genes such as *IFIT1* and *ISG15* (**Supp Fig. 3J,K**). These findings suggest that dopamine selectively engages NF-κB-dependent signaling downstream of cGAS-STING, rather than triggering a broader interferon-associated response, consistent with the lack of broader cytokine response shown in the Mesoscale multiplex assay. To confirm that cGAS-STING signaling contributes to dopamine-induced increases in IL-6 secretion, hMDMs were treated with dopamine following pharmacological inhibition (1 hr) of either cGAS (G140, 1uM)^48^ or STING (H-151, 1uM)^49^. Inhibition of either pathway significantly attenuated dopamine-induced IL-6 secretion, as measured by AlphaLISA (**Fig. 3G-I**), demonstrating that cGAS-STING activation is required for dopamine-driven increases in IL-6 production in hMDM.

### Dopamine triggers mitochondrial dysfunction and mtDNA release to activate cGAS-STING signaling

Activation of the cGAS-STING pathway is initiated by exposure to double stranded DNA (dsDNA) in the cytoplasm^42^. Mitochondrial DNA (mtDNA) is a circular, dsDNA that is normally confined within mitochondria but can be released into the cytoplasm following mitochondrial stress or damage^50,51^. In the cytoplasm, it acts as a potent endogenous danger signal, initiating multiple inflammatory pathways, including cGAS-STING. To determine if dopamine damages mitochondria and induces mtDNA release in the cytoplasm, we isolated DNA from the cytoplasmic fraction of dopamine-treated hMDMs (1 hr and 6 hrs) and assessed mtDNA levels via qPCR. After 6 hrs, there was a significant increase in mtDNA abundance, as measured by Cytochrome C transcript, within the cytoplasmic fraction. This was significantly higher than the levels of cytoplasmic mtDNA after 1 hr of dopamine exposure or compared with 1 or 6 hrs of vehicle treatment (**Fig. 4A**). Assessment of cell numbers and viability using cell counts showed no changes with dopamine treatment, indicating that increased mtDNA was not due to cell death (**Supp Fig. 3L**). To determine whether the dopamine mediated increase in mtDNA mediates the effects of dopamine on cGAS-STING activity, hMDM were treated with ethidium bromide (EtBr, 10 μM) to deplete cellular mtDNA^52,53^. Decreasing cellular mtDNA with EtBr abrogated dopamine-induced activation of the cGAS-STING pathway (**Fig. 4B-D**) and significantly reduced IL-6 secretion (**Fig. 4E**), indicating that mtDNA is necessary for dopamine-driven inflammatory signaling.

**Figure 4.**
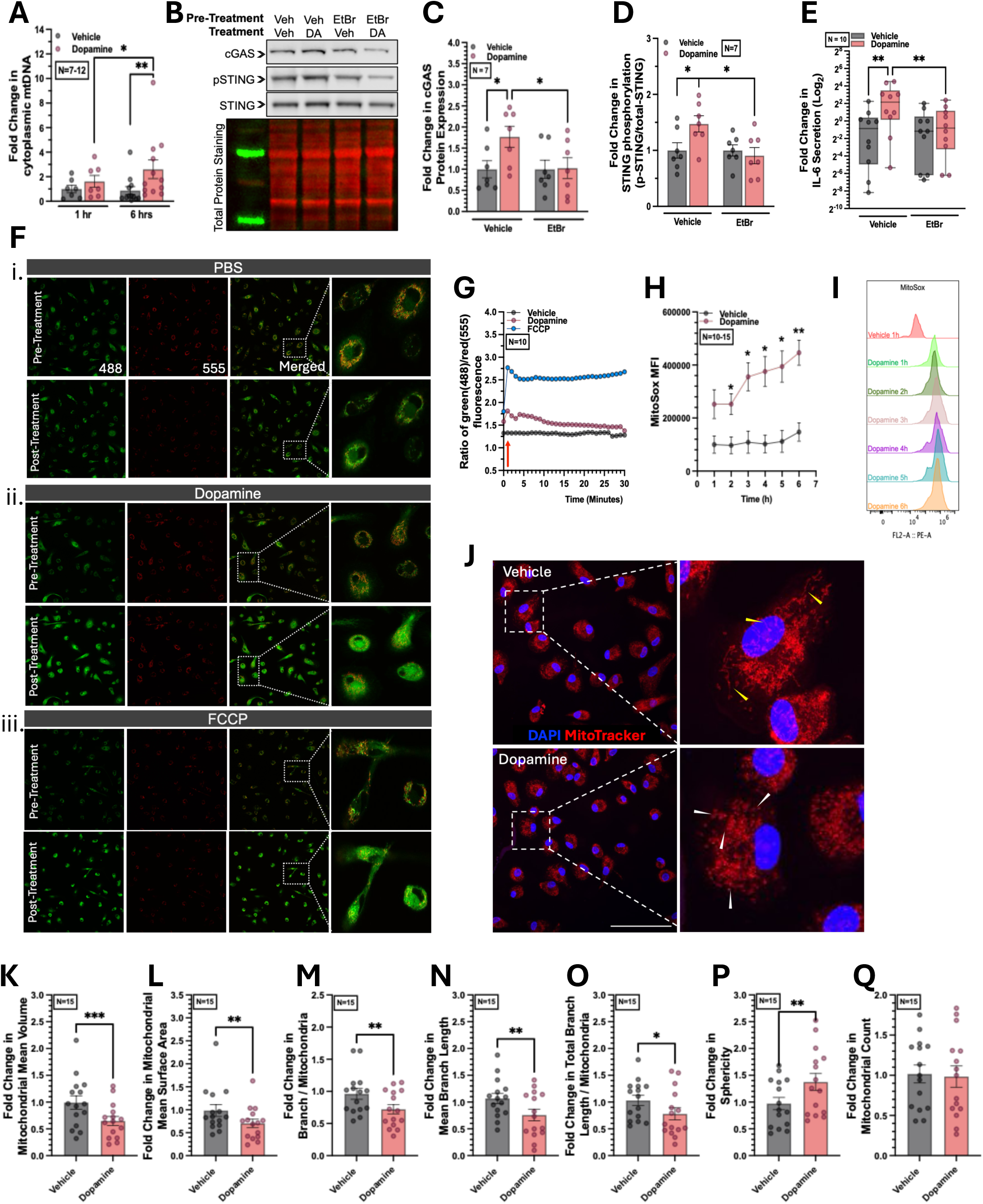
Dopamine triggers mitochondrial dysfunction and mtDNA release to activate cGAS-STING signaling. (A) Cells were treated with vehicle or dopamine and collected at 1 hr or 6 hrs post-treatment. Following mitochondrial and cytosolic fractionation, DNA was isolated from the cytoplasmic fraction, and qPCR was performed to measure cytochrome c mitochondrial DNA levels relative to 18S, indicating mtDNA release into the cytoplasm. (Biological replicates: N=7-12). Data are presented as mean±SEM. (B-D) hMDMs were pre-treated with Vehicle or EtBr (10 μM, t = −1 h) to deplete mitochondrial DNA, followed by treatment with vehicle or dopamine for 6 hrs. Cells were lysed and assessed for cGAS protein expression as well as phosphorylation levels of STING by western blotting. (Biological replicates: N=7). Data are presented as mean±SEM. (E) Supernatants from experiments in panel B-D were collected 24 hrs post-treatment, and IL-6 secretion was measured using AlphaLISA. (Biological replicates: N=10). Data are presented as box and whisker plot (min to max). (F,G) Using confocal live imaging, JC-1 stained cells were imaged every 1 minute, for 30 minutes. Following the first image, cells were treated with vehicle (PBS) (C-i), dopamine (C-ii), or a positive control (FCCP, 1.5 μM) (C-iii) (t:1 min). The scale bar represents 20 μm. The ratio of green fluorescence (488 nm) to red fluorescence (555 nm) was calculated as an indicator of mitochondrial membrane depolarization, (Biological replicates: N=10). Data are shown as mean of response in each condition at each specific timepoint. (H,I) Cells were treated with vehicle or dopamine in a time-course experiment (1 - 6 hrs) and then stained with MitoSOX. Using flow cytometry, the mean fluorescence intensity (MFI) of MitoSOX staining was measured in the live singlet population to assess mitochondrial superoxide production. (Biological replicates: N=10-15). Data are presented as mean±SEM and representive histogram shows MitoSox MFI overtime. (J-Q) Cells were treated with vehicle or dopamine for 6 hrs and stained with MitoTracker Orange CMTMRos. Following fixation, nuclei were stained with DAPI, and cells were imaged using confocal microscopy (30× objective) with 3D acquisition at a step size of 0.15 μm. Yellow arrows represent elongated mitochondria, while white arrows represent round-shape mitochondria (fission). The scale bar represents 20 μm. Images were processed using ImageJ, and representative images are shown as z-projections for each condition. Mitochondrial morphology analysis was performed using the Mitochondria Analyzer plug-in for FIJI, (Biological replicates: N=15). Data are presented as mean±SEM. Paired comparisons were analyzed using a paired *t*-test (non-parametric: Wilcoxon test) in panel H and K-Q or paired one-way ANOVA (non-parametric: Friedman test) in panel C-E, or unpaired one-way ANOVA (non-parametric: Kruskal Wallis test) in panel A. Significance levels are indicated as follows: *p* < 0.05 (*), *p* < 0.01 (**), *p* < 0.001 (***), *p* < 0.0001 (****).

Release of mtDNA is indicative of mitochondrial damage, so we evaluated the impact of dopamine on mitochondrial health, starting with assessment of mitochondrial membrane potential (ΔΨm) using JC-1 staining. Dopamine treatment induced a rapid depolarization of the mitochondrial membrane immediately following exposure, as shown by the increase in the green to red fluorescence ratio, corresponding to an increase in monomeric JC-1 (low membrane potential) relative to J-aggregates (high membrane potential). Dopamine-induced depolarization was transient, with membrane potential largely recovering within 10 -15 min after treatment. This contrasts with the positive control FCCP (1.5 µM), which caused rapid and sustained mitochondrial depolarization (**Fig. 4F,G**). Transient mitochondrial depolarization, or “flickering” can precede significant mitochondrial dysfunction and is often associated with rapid bursts in the production of mitochondrial reactive oxygen species such as mitochondrial superoxide (MitoSox)^54,55^. Indeed, flow cytometry showed that dopamine treatment induced a sustained, significant increase in mitochondrial superoxide levels, with maximal effects observed at 6 hr post-stimulation (**Fig. 4H,I**). In addition to depolarizing mitochondria and increasing MitoSox levels, dopamine also induced changes in mitochondrial morphology. Confocal imaging of hMDM stained with MitoTracker™ Orange CMTMRos showed that dopamine resulted in pronounced mitochondrial fragmentation consistent with increased mitochondrial fission (**Fig. 4J**). Morphometric analyses revealed that dopamine treatment significantly reduced mitochondrial mean volume (**Fig. 4K**), surface area (**Fig. 4L**), branch length and count (**Fig. 4M-O**) and increased mitochondrial sphericity, resulting in rounder, less elongated mitochondria (**Fig. 4P**). As total mitochondrial counts were not significantly altered, this likely reflects a balance between increased fission and ongoing mitochondrial damage and rupture leading to mtDNA release (**Fig. 4Q**).

### DRP1-Mediated Mitochondrial Fission Drives mtDNA Release and cGAS-STING Activation in Response to Dopamine

Mitochondrial fission is regulated by dynamin-related protein 1 (DRP1), a cytosolic GTPase that mediates mitochondrial fission upon phosphorylation and recruitment to the mitochondrial outer membrane^56^. Assessing the impact of dopamine on DRP-1 over time demonstrated that dopamine treatment induced robust DRP1 phosphorylation (Ser616) beginning at 1 hr. This was sustained and increased by 6 hrs (**Fig. 5 A,B**), suggesting that dopamine-induced DRP-1 phosphorylation and therefore mitochondrial fission may drive the effects of dopamine in hMDM. To determine if this change in DRP1-mediated mitochondrial fission was necessary for dopamine-mediated cGAS-STING activation and IL-6 secretion, hMDM were pre-treated with Mdivi-1 (40 μM, selective DRP-1 inhibitor) for 1 hr^57,58^, then stimulated with dopamine. Mdivi-1 eliminated the dopamine induced alterations in mitochondrial morphology and reduced phosphorylation of DRP1 (**Fig. 5C-J**). Inhibition of DRP1 activity further revealed that mitochondrial fission is required for downstream signaling, as Mdivi-1 treatment attenuated dopamine-induced upregulation of cGAS and phosphorylation of STING (**Fig. 5K-M**). Consistent with this, blockade of DRP1-mediated fission significantly reduced IL-6 secretion in response to dopamine stimulation (**Fig. 5N**). Moreover, Mdivi-1 treatment mitigated dopamine-induced mitochondrial dysfunction, demonstrated by decreased MitoSox production (**Fig. 5O**) and reduced release of mtDNA into the cytoplasm (**Fig. 5P**). These data show that DRP1-dependent mitochondrial fission is a central drive of dopamine-induced mitochondrial dysfunction, and a critical link to activation of the cGAS-STING pathway and IL-6 production.

**Figure 5.**
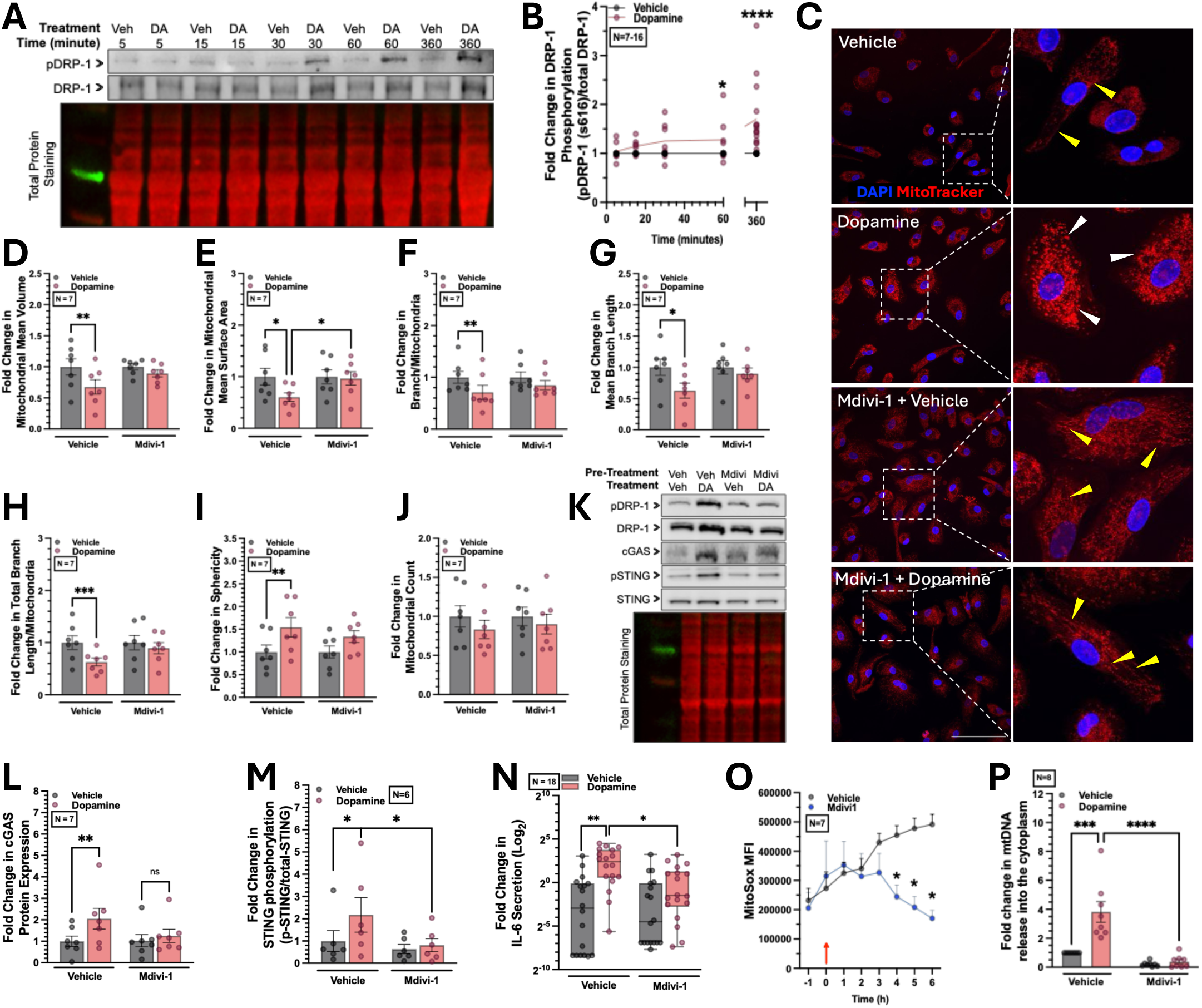
DRP1-Mediated Mitochondrial Fission Drives mtDNA Release and cGAS-STING Activation in Response to Dopamine. (A,B) Cells were treated with vehicle or dopamine in a time-course experiment (5, 15, 30, 60, and 360 minutes). Phosphorylation levels of DRP1 (ser616) were measured by western blotting and quantified as the ratio of phosphorylated DRP1 to total DRP1. The data suggest that dopamine induces DRP1 phosphorylation over time, with maximal responses observed at 6 hrs post-treatment (Biological replicates: N=7-16). Individual values are shown with line demonstrating mean. (C) hMDMs were pre-treated with Vehicle or Mdivi-1 (40 μM, t = −1 h) to inhibit DRP1-mediated mitochondrial fission, followed by treatment with vehicle or dopamine for 6 hrs. Cells were stained with MitoTracker Orange CMTMRos. Following fixation, nuclei were stained with DAPI, and cells were imaged using confocal microscopy (30× objective) with 3D acquisition at a step size of 0.15 μm. The scale bar represents 20 μm. Yellow arrows represent elongated mitochondria, while white arrows represent round-shape mitochondria (fission). Images were processed using ImageJ, and representative images are shown as z-projections. (D-J) Mitochondrial morphology parameters were analysis using the Mitochondria Analyzer plug-in for FIJI. (Biological replicate: N=7). Data are presented as mean±SEM. (K-M) hMDMs were pre-treated with Vehicle or Mdivi (40 μM, t = −1 h) to inhibit DRP-1 mediated fission, followed by treatment with vehicle or dopamine for 6 hrs. Cells were lysed and assessed for cGAS protein expression as well as phosphorylation levels of STING by western blotting (Biological replicates: N=6 for cGAS or N=7 for pSTING). Data are presented as mean±SEM. (N) Supernatants from experiments in panel K-M were collected 24 hrs post-treatment, and IL-6 secretion was measured using AlphaLISA (Biological replicates: N=18). Data are presented as box and whisker plot (min to max). (O) Cells were pretreated with Mdivi-1 (t=-1) and were stained with MitoSOX, and MitoSox production was assessed by flow cytometry for cells treated with dopamine at t=0 and collected at different timepoints shown in x-axis. The MFI of MitoSOX staining was measured in the live singlet population. Red arrow represents dopamine treatment at t=0 (Biological replicates: N=7). Data are presented as mean±SEM. (P) Cells were treated similar to panel K=M and were followed by mitochondrial and cytosolic fractionation. DNA was isolated from the cytoplasmic fraction, and qPCR was performed to measure mtDNA levels relative to 18S, indicating mtDNA release into the cytoplasm. (Biological replicates: N=8). Data are presented as mean±SEM. Paired comparisons were analyzed using a paired t-test (non-parametric: Wilcoxon test) in panel B and O, or paired one-way ANOVA (non-parametric: Friedman test) in panel D-I, L-N and P. Significance levels are indicated as follows: p < 0.05 (*), p < 0.01 (**), p < 0.001 (***), p < 0.0001 (****).

### Dopamine induces a metabolic shift from oxidative phosphorylation to glycolysis

Elevated production of MitoSOX due to excessive mitochondrial fission is often associated with impaired electron transport chain (ETC) function, reduced oxidative phosphorylation (OXPHOS), and altered ATP production. Cells can compensate through glycolytic shift, i.e. a metabolic shift toward aerobic glycolysis^59,60^, which many studies have linked to increased activation in innate immune cells such as macrophages^61,62^. To determine whether dopamine-induced mitochondrial alterations lead to glycolytic shift, we assessed mitochondrial respiration and glycolytic activity at 1 hr and 6 hr post-dopamine treatment. Analysis using Seahorse ATP rate assay showed that 1 hr of dopamine exposure reduced baseline oxygen consumption rate (OCR), while extracellular acidification rate (ECAR) remained unchanged (**Fig. 6A,B**). Despite this reduction in basal respiration, total ATP production and the relative contribution of OXPHOS versus glycolysis to ATP generation were not significantly altered (**Fig. 6C,D**). In contrast, at 6 hrs post-dopamine treatment, OCR levels returned to baseline, but ECAR was significantly increased (**Fig. 6E,F**), indicating enhanced glycolytic activity. While total ATP production remained comparable to vehicle-treated cells, a significantly greater proportion of ATP was generated through glycolysis, consistent with a metabolic shift away from OXPHOS (**Fig. 6G,H**). These findings suggest dopamine induces an early but limited impact on cellular energetics, followed by a sustained metabolic shift towards glycolysis within the mitochondrial population, despite partial restoration of mitochondrial respiration. This was further explored using a Seahorse Mito Stress assay (**Fig. 6I**). This assay confirmed the prior findings, demonstrating that dopamine induces early (1 hr) alterations in mitochondrial respiration that largely returns to baseline levels at later (6 hr) time points. These effects were most evident in basal and maximal respiration, whereas proton leak, spare respiratory capacity, coupling efficiency, and ATP-linked respiration remained comparable between groups (**Fig. 6J-O**).

**Figure 6.**
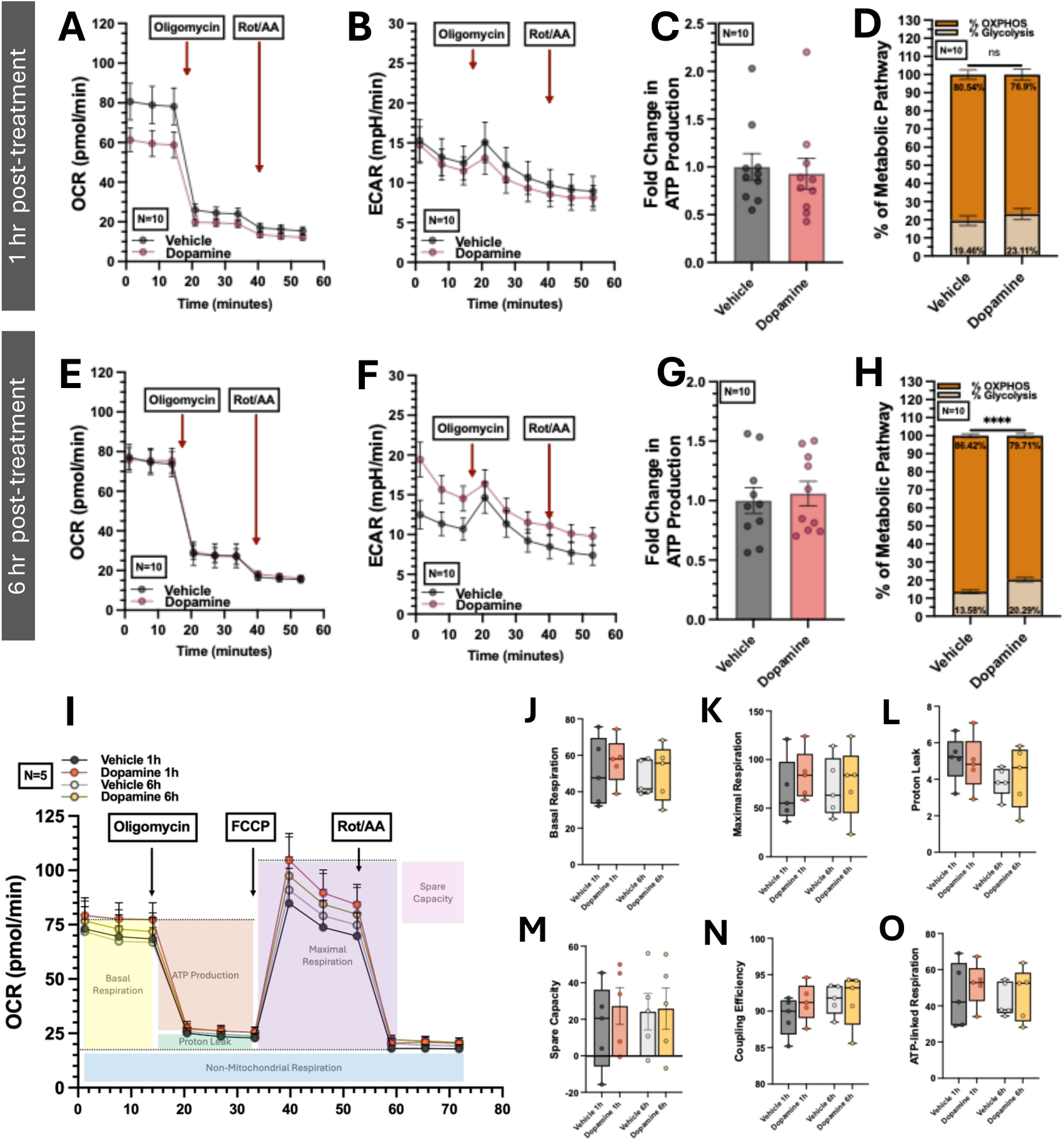
Dopamine induces an imbalance in cellular respiration, with an increased shift toward glycolysis over-time. (A-D) hMDMs were treated with vehicle or dopamine for 1 hr and analyzed using the ATP Rate Assay on an Agilent Seahorse XF 96-well analyzer. During the assay, cells were sequentially treated with oligomycin and rotenone/antimycin A to inhibit ATP synthase (Complex V) and Complex I/III, respectively. Data were analyzed using the Seahorse Analytics software (Biological replicate: N=10). Data are presented as mean±SEM. (E-H) hMDMs were treated with vehicle or dopamine for 6 hrs and analyzed using the ATP Rate Assay on an Agilent Seahorse XF 96-well analyzer. During the assay, cells were sequentially treated with oligomycin and rotenone/antimycin A to inhibit ATP synthase (Complex V) and Complex I/III, respectively. Data were analyzed using the Seahorse Analytics software (Biological replicate: N=10). Data are presented as mean±SEM. (I-O) To further assess the time-dependent differences (1 hr vs 6 hrs) following dopamine treatment, a Mito Stress Assay was performed using the Agilent Seahorse XF 96-well analyzer. During the assay, cells were sequentially treated with oligomycin, FCCP, and rotenone/antimycin A to disrupt ATP synthase (Complex V), the proton gradient, and Complex I/III, respectively. Data were analyzed using the Seahorse Analytics software (Biological replicate: N=5). Data are presented as mean±SEM in panel I and as as box and whisker plot (min to max) in panel J-O. Paired comparisons were analyzed using a paired *t*-test (non-parametric: Wilcoxon test) in panel H. Significance levels are indicated as follows: *p* < 0.0001 (****).

These data suggest that the early dopamine induced perturbations in the mitochondrial compartment (mitochondrial membrane potential depolarization) leave a sustained impact on mitochondrial metabolism. These types of perturbations in mitochondrial populations often result in mitophagy, or selective mitochondrial autophagy, to remove defective mitochondria^63,64^. PINK1 and PARKIN are central mediators of ubiquitin-dependent mitophagy, while BNIP3/NIX and LC3B are involved in receptor-mediated mitochondrial clearance and autophagosome formation, respectively^65–67^. To determine whether dopamine induced mitochondrial dysfunction was associated with increase in mitophagy, the activity of the key mitophagy regulators PINK1, PARKIN, BNIP3/NIX and LC3B were examined at 1 hr and 6 hrs post-dopamine treatment. Dopamine induced a modest, non-significant increase in PINK1 expression at 1 hr, with no detectable changes in PARKIN, BNIP3/NIX, or LC3B levels (**Supp Fig. 4A-E**), suggesting that neither pathway was active. To further assess mitophagic flux, we examined the interaction of mitochondria with lysosomes by evaluating the colocalization of TOMM20 (mitochondrial outer membrane marker) and LAMP1 (lysosomal marker). Dopamine treatment did not increase TOMM20-LAMP1 colocalization at either time point, indicating that mitophagy is not induced by dopamine (**Supp Fig. 4F-I**). These findings suggest that dopamine-induced mitochondrial alterations may occur in the absence of efficient mitochondrial clearance, potentially allowing the accumulation of damaged mitochondria.

## Discussion

Dopamine is increasingly viewed as an immunomodulatory factor, although studies show that dopamine can have both pro- and anti-inflammatory effects depending on the experimental paradigm^4^. Our prior studies showed that dopamine enhances human macrophage immune activity, including the release of inflammatory cytokines such as IL-6 and IL-1β; however, the underlying mechanisms remained undefined. Here, we show for the first time that dopamine promotes macrophage inflammatory responses, specifically IL-6 secretion, through disruption of mitochondrial homeostasis via excessive DRP1-dependent mitochondrial fission. This process leads to the cytoplasmic release of mtDNA, which functions as a danger-associated molecular pattern (DAMP) to activate cGAS-STING pathway. Activation of this pathway enhances NF-κB signaling, thereby driving IL-6 production. Prior studies have established mitochondrial metabolism as a key regulator of macrophage activation and inflammatory function^68,69^, and metabolic shifts toward aerobic glycolysis support the production of pro-inflammatory cytokines, including IL-6^69^. This reprogramming can arise from mitochondrial dysfunction, including increased ROS, disruption of mitochondrial fission/fusion balance, and impaired activity of the ETC, all of which reinforce glycolytic metabolism^27^. While there is a growing awareness of a link between neurotransmission in mitochondrial dysfunction^70–72^, currently there is no defined connection between dopamine and the regulation of mitochondrial metabolism.

In this study, dopamine-induced mitochondrial alterations appear to follow a progressive and self-amplifying pattern. Early responses include transient depolarization of the mitochondrial membrane potential, which appears reversible at early time points. However, downstream events such as DRP1-mediated mitochondrial fission and MitoSox production seem to progressively increase over time. The transient depolarization phase may represent a period of mitochondrial “flickering,” during which mitochondrial dysfunction remains reversible^73^. As mitochondrial stress and excessive fission persist, however, these early alterations appear to transition into sustained mitochondrial impairment. This feed-forward mitochondrial damage likely contributes to progressively stronger inflammatory outputs over time. At early time points cytoplasmic mtDNA levels show only a trend toward an increase with dopamine. In contrast, later time points show an amplification robust mitochondrial fission and significant changes in mitochondrial morphology, elevated mitochondrial ROS production, a metabolic shift toward glycolysis and robust increase in mtDNA release into the cytoplasm.

Despite these alterations, total mitochondrial counts remain relatively unchanged, which might be partially due to dysregulated mitophagy. This can be additionally explained by the transient depolarization of mitochondrial membrane (∼10-15 min), which may not be sufficient to fully activate the mitophagy pathway^74^. Indeed, impaired mitophagy has been reported to promote the accumulation and cytoplasmic release of mtDNA, subsequently activating innate immune signaling pathways^75^. This supports the activity seen in response to dopamine in this study. The metabolic consequences of these mitochondrial alterations further support this model. At early time points, dopamine reduces oxygen consumption rates while maintaining extracellular acidification rates, suggesting an early impairment in mitochondrial respiration without immediate metabolic compensation. At later time points, however, extracellular acidification rates increase significantly while overall ATP production remains stable.

Release of mtDNA into the cytoplasm is a key consequence of mitochondrial damage, as cytoplasmic DNA can activate multiple innate immune sensing pathways, including TLR9 ^76^, cGAS-STING ^77^, and ZBP1 pathways^78^, as well as the NLRP3^79^ and AIM2 inflammasomes^80^. Our data indicate that dopamine activates the cGAS-STING pathway, in which cGAS detects cytosolic DNA and synthesizes the second messenger 2′3′-cGAMP, which activates STING. Activated STING becomes phosphorylated at the endoplasmic reticulum, transfers to the Golgi, and recruits downstream signaling molecules such as TBK1. This typically leads to phosphorylation of IRF3 and activation of type I interferon responses, and/or activation of NF-κB^44^. Our data demonstrate that dopamine induces activation of the cGAS-STING pathway over time, correlating with mtDNA accumulation observed in the cytoplasm. Interestingly, we did not observe activation or phosphorylation of TBK1 in this system. Previous studies have shown that TBK1 activation is strongly associated with IRF3-dependent transcriptional responses, while NF-κB activation in myeloid cells can occur independently of TBK1 and IRF3 signaling. These findings support a model in which dopamine-induced mtDNA release activates cGAS-STING signaling that preferentially drives NF-κB-mediated inflammatory responses rather than canonical TBK1-IRF3 interferon signaling^44,81–85^. Confirming the role of dopamine-induced cGAS-STING activation in driving IL-6 secretion, pharmacological inhibition strategies targeting NF-kB, cGAS, and STING all significantly reduced dopamine-induced IL-6 secretion. Notably, these data show that only dopamine, rather than its metabolic products (DOPAC, HVA), precursors (L-DOPA) or other catecholamines (serotonin, norepinephrine), induces these effects on IL-6 production in hMDMs. Although macrophages express enzymes capable of dopamine synthesis and metabolism^86^, the inability of these metabolites to induce IL-6 secretion emphasizes the importance of dopamine itself as the primary stimulus in this pathway. This mechanism may, at least in part, contribute to the elevated IL-6 levels observed in the serum of individuals using substances of misuse, including cocaine, which are associated with increased dopamine levels in the body^87^.

These data support prior studies^9,10,17^ showing dopamine is a pro-inflammatory stimulus for human macrophages. The discrepancy between rodent and human dopamine-mediated responses may arise from several factors. Differences in immune signaling machinery, the presence or absence of certain regulatory proteins, and species-specific immune responses may all contribute to these contrasting outcomes^60–62^. However, differential expression levels of dopamine receptors may also play a central role in determining whether dopamine signaling leads to pro- or anti-inflammatory responses^91–93^. Notably, the variability observed between donors in this cohort is characteristic of primary human cells and reflects physiological differences in dopamine receptor expression and immune responsiveness among donors. Despite that variability, the data here show a clear correlation between dopamine receptor expression levels, specifically expression of DRD1, and dopamine-driven IL-6 release, suggesting that dopamine receptor expression regulates dopamine-mediated inflammatory responses in human macrophages. Our prior work in human macrophages demonstrated a similar correlation between dopamine receptor expression and dopamine-induced IL-1β production, supporting the importance of receptor expression levels in determining inflammatory outcomes. Studies in rodents have also shown that expression of DRD5 regulates CD4+ T-cell inflammation in the gut^94^, and that dopamine can induce IL-6 secretion through D1-like receptors and promote morphological changes in astrocytic cells, although additional receptors such as adrenergic receptors may also contribute to the response^95^. These findings highlight the complexity of dopamine signaling and suggest that receptor expression patterns may influence inflammatory outputs.

The specific connection between dopamine receptor signaling and the early mitochondrial effects of dopamine exposure remain unclear. Extensive analysis of signaling pathways demonstrated that dopamine is not activating MAP kinases, AKT or the JAK/STAT pathway on a time scale that would impact this process. Importantly, the mitochondrial responses to dopamine, specifically the rapid depolarization of the mitochondrial membrane, begin within seconds of dopamine exposure, supporting the involvement of signaling mechanisms that operate independently of slower transcriptional processes. Dopamine receptors are G-protein coupled receptors (GPCRs) that classically signal through distinct G-protein subunits, leading to modulation of cyclic AMP (cAMP) levels or activation of diacylglycerol (DAG) and inositol trisphosphate (IP_3_) signaling pathways^96,97^. However, prior studies have shown that dopamine receptors do not induce cAMP production in dopamine-induced responses in hMDM^15^. This suggests that calcium signaling, which is induced by dopamine receptors in hMDM^15^, could be a primary mediator of this process, as numerous studies have linked mitochondrial damage to calcium signaling^98,99^. Additionally, dopamine receptors and organic cation transporters (OCTs) have been reported to localize to intracellular organelles^100,101^, raising the possibility that direct mitochondrial dopamine signaling may contribute to the observed mitochondrial responses. And while our data show that dopamine receptor expression is clearly associated with the dopamine mediated effect on IL-6, dopamine receptors may act in concert with other dopaminergic machinery, such as the dopamine transporter (DAT). DAT is also expressed on macrophages, and has been shown to regulate hMDM IL-6 release and can mediate intracellular dopamine signaling^102^. Overall, these data define a previously unrecognized mechanism linking dopamine to mitochondrial dysregulation and innate immune activity, specifically through changes in mitochondrial metabolism and activation of the cGAS-STING pathway. These findings further support the role of dopamine as an important, unrecognized immunomodulatory factor, and suggest that dopamine mediates its immunoregulatory effects via unique signaling pathways in human primary macrophages.

### Limitations of the study

Dopamine presents several technical limitations in experimental settings due to its instability, short half-life in solution, and high sensitivity to light. These properties limit the ability to perform extended functional assays in vitro, including prolonged or repeated measurements for cellular respiration, as well as long-term live-cell imaging approaches. Additionally, the use of primary human macrophages introduces both strengths and limitations to this study. While donor-to-donor heterogeneity and variable cellular responses enhance the translational relevance of our findings, they also increase experimental complexity and necessitate larger sample sizes (N) to achieve adequate statistical power compared to studies using clonal cell lines. This work is specifically focused on human myeloid cells, and it is critical to note that dopamine receptor expression and signaling profiles differ significantly between humans and rodent or immortalized cell systems, limiting the direct translatability of those models. Furthermore, the use of primary human macrophages limits the application of genetic manipulation approaches, such as knockdown or knockout strategies, which could provide stronger mechanistic insights. These techniques remain technically challenging in primary cells due to their limited proliferative capacity and sensitivity to extended culture conditions.

### Resource availability

#### Lead Contact

Request for further information and resources should be directed to the lead contact, Peter J. Gaskill

#### Material Availability

This study did not generate new unique reagents.

#### Data and code availability

The mass spectrometry data generated in this study have been deposited in the ProteomeXchange Consortium through the MassIVE repository under the dataset identifier PXD077267. Code for the principal component analysis and cluster analysis is posted to Zenodo at this DOI: https://doi.org/10.5281/zenodo.19670069

## Acknowledgments

The authors thank all donors who contributed PBMCs to this study. We also thank members of the Gaskill Lab for their contributions, as well as for valuable feedback and discussions. This work was supported by the Schlumberger Faculty for the Future Fellowship (MD); the National Institute of Allergy and Infectious Diseases (F30AI179472 to BC); the National Institute on Drug Abuse (R01DA057337, R33DA05850103, R01DA062277 to PJG; R61DA061825 to JGJ; U01 DA053624 to H.F); the National Institute of Mental Health (K01MH132466 to SMM; P30 MH092177 funding WD [CNHC, CTRSC, Drexel Component; PI: BW]); the National Institute on Aging (R01AG081929 to JGJ); the Congressionally Directed Medical Research Programs (ME220134, ME240242 to JGJ); and the WW Smith Charitable Trust (to SMM and JGJ). H.-Y.T. is supported by R50CA221838. Funding for the proteomics facility at The Wistar Institute was provided by the Cancer Center Support Grant (P30CA010815).

## Author contributions

Conceptualization: M.D., B.C., K.S., J.G.J., P.J.G.

Data Curation: M.D., B.C., E.C., L.S., A.M.F.,

Formal Analysis: M.D., A.M.F., H-Y. T., W.D., B.C., E.C., A.A.

Funding acquisition: M.D., B.C., S.M.M., J.G.J., W.D., H.F., P.J.G.

Investigation: M.D., B.C., E.C., A.A., L.S., T.K., S.K., J.M., A.M.F., T.B.

Methodology; M.D., B.C., P.J.G.

Project Administration: M.D, P.J.G.

Resources: M.D., B.C., E.C., J.M., T.K., A.M.F. T.B., H-Y. T.

Software: M.D., A.M.F., W.D.

Supervision: P.J.G.

Validation: M.D., W.D., A.F., P.J.G.

Visualization: M.D., W.D., P.J.G.

Writing - original draft: M.D.

Writing - review & editing: M.D., B.C., E.C., J.G.J., H-Y. T., A.M.F., S.M.M., W.D., H.F., P.J.G.

## Supplemental information titles and legends

Document S1: Supplemental Figures 1-4, Supplementary Tables 1 and 2

---

**Supplementary Figure 1.**
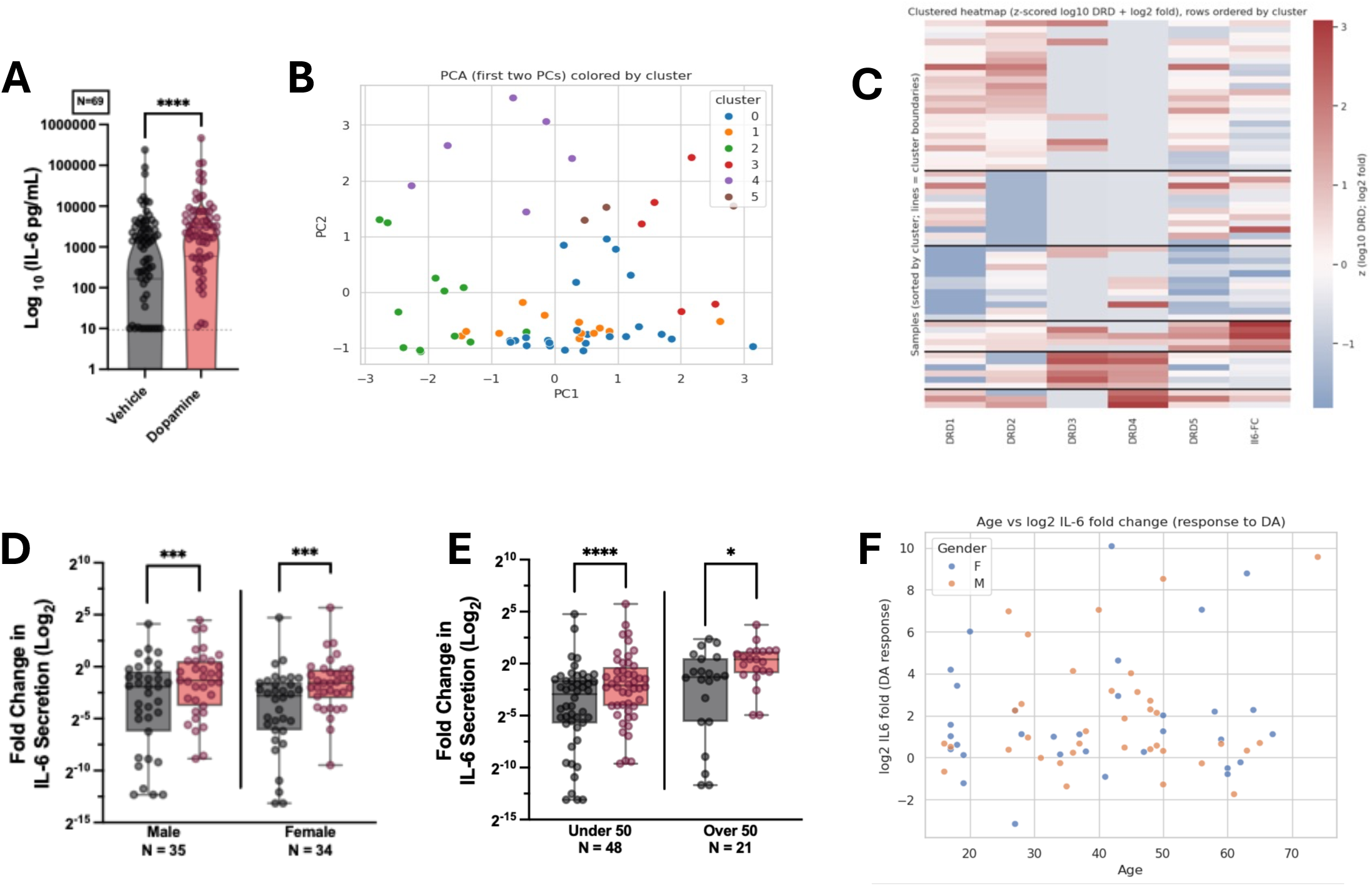
Multivariate and demographic analyses reveal donor heterogeneity in dopamine-stimulated hMDMs. (A) Dopamine treatment induces a significant increase in IL-6 secretion 24 hours post-treatment, as measured by AlphaLISA and data reported as pg/mL values. The dashed line indicates the limit of detection (Biological replicate: N=69). Data are presented as volcano plot with individual donors shown on the graph. (B) PCA biplot of PC1 vs. PC2 for the 62 complete-case donors, colored by cluster assignment (K = 6). PC1 and PC2 capture 57.2% of total variance. The two smallest clusters (C4, n = 2; C5, n = 4) occupy peripheral positions in this space (Biological replicate: N=62). Data are presented as scatter plot. (C) Cluster-sorted sample × feature heatmap. Rows are donors (n = 62) sorted by cluster assignment (K = 6); columns are the six panel features; color encodes z-score after log₁₀ (DRD) / log₂ (fold) transforms. Horizontal black lines mark cluster boundaries. Row labels are suppressed for clarity (Biological replicate: N=62). (D,E) Donors were stratified based on the gender (Male vs. Female) and age (under- or over-50) and IL-6 secretion levels (AlphaLISA) were compared between groups (Biological replicate: N=69). Data are presented as box and whisker plot (min to max). (F) Bivariate relationship between donor age and log₂ IL-6 fold change (in response to dopamine). Each point is one donor; point color encodes gender. (Biological replicate: N=62). Data are presented as scatter plot. Paired comparisons were analyzed using a paired *t*-test (non-parametric: Wilcoxon test) in panel A. Comparisons between Gender/Age groups were performed using an unpaired *t*-test (Mann-Whitney test) in panel D and E. Significance levels are indicated as follows: *p* < 0.05 (*), *p* < 0.01 (**), *p* < 0.001 (***), and *p* < 0.0001 (****).

**Supplementary Figure 2.**
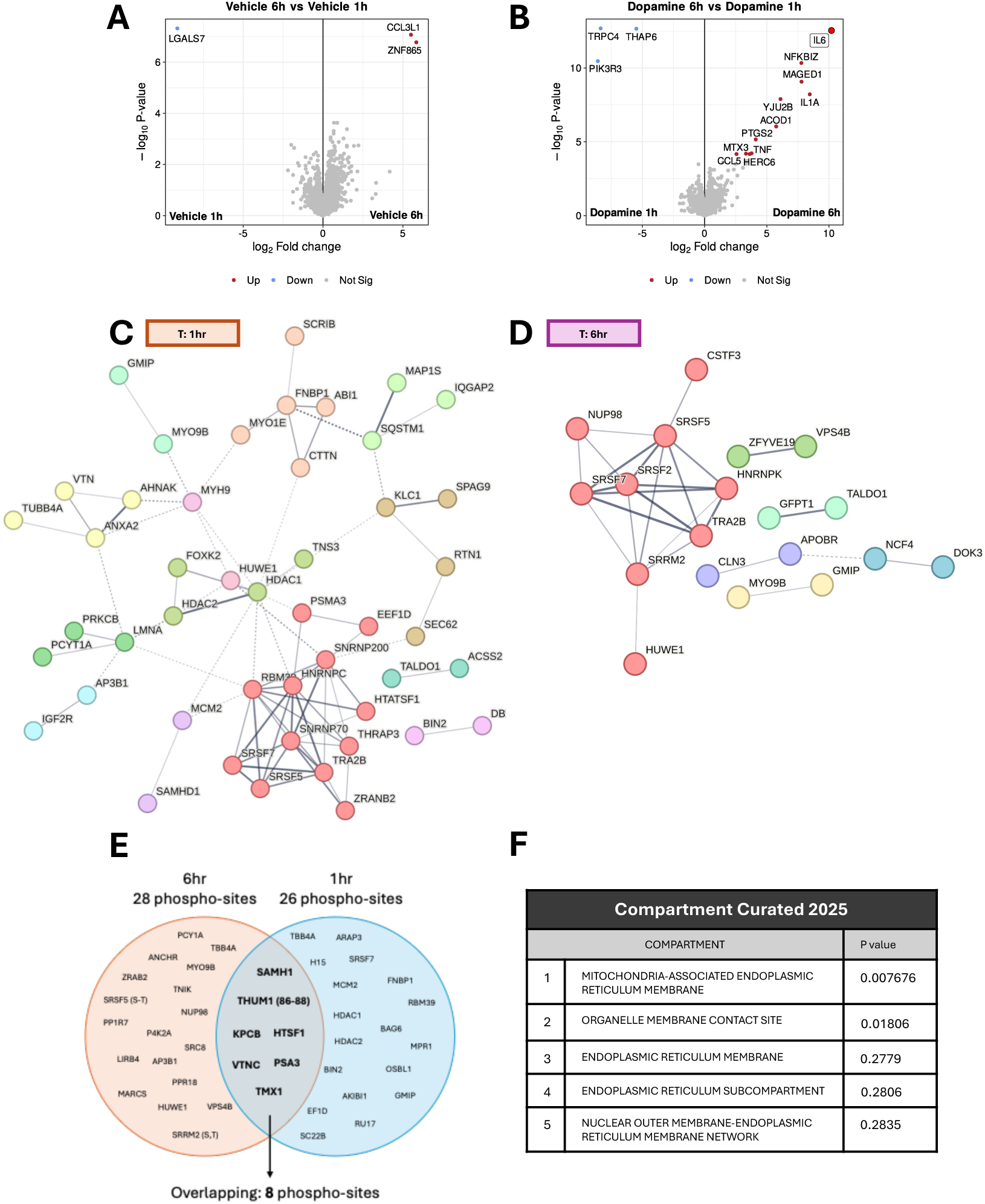
Dopamine Reprograms Proteomic and Phospho-Signaling Landscapes. (A,B) hMDMs from five distinct donor groups were treated with vehicle or dopamine (1 μM) and collected at 1 hr or 6 hrs post-treatment. Proteins with significantly altered abundances between treatment groups, to compare the over-time alterations, are shown in volcano plots (Biological replicate: N=5). (C,D) STRING interaction enrichment analysis was performed on proteins with ≥2-fold increased phosphorylation, highlighting biological interactions between proteins, within each timepoint group. Edges represent protein-protein associations, with edge confidence ranging from low (0.150), medium (0.400), high (0.700) and highest (0.900), represented by line thickness. Node colors indicate clustering based on Markov Cluster (MCL) algorithm. (E,F) Proteins with significantly increased phosphorylation following dopamine treatment are shown in a Venn diagram, with 6 hr treatment (orange) and 1 hr treatment (blue). Overlapping proteins are listed in the center, and these were used for compartment-curated analysis, with associated compartments and their p-values indicated.

**Supplementary Figure 3.**
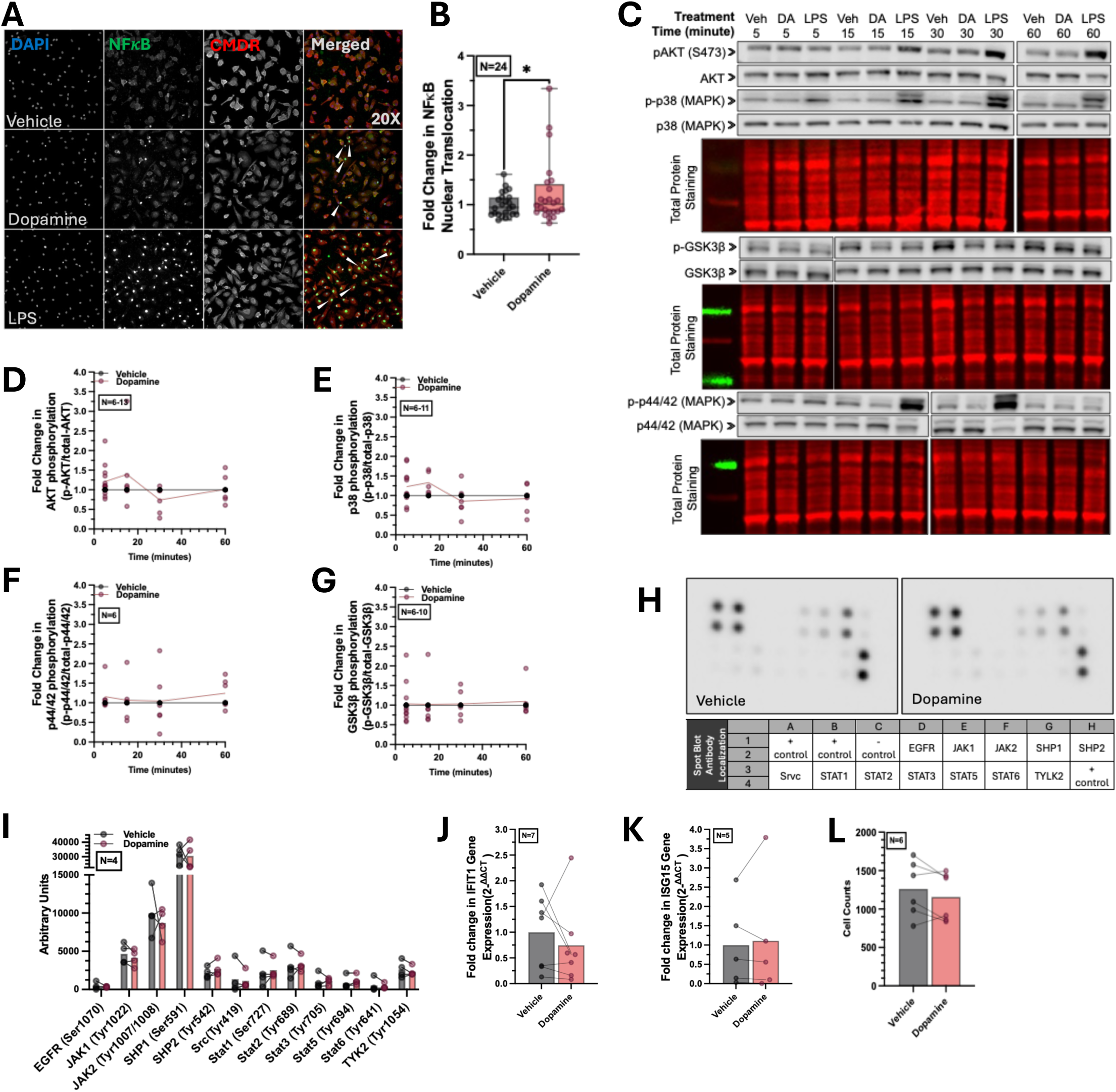
Dopamine Activates NF-κB Independent of Canonical Signaling Pathways. (A,B) High-content imaging shows that dopamine induces a modest increase in the nuclear translocation of p-NFκB at 1 hr post-treatment. This was quantified by calculating the ratio of p-NFκB intensity (green) in the nucleus (DAPI, blue) to that in the cytoplasm. The cytoplasmic region was defined using CellMask Deep Red (CMDR, red), and the cytoplasmic signal was calculated by subtracting the nuclear region (DAPI) from the CMDR mask. Cells with high nuclear translocation values are indicated by white arrows. LPS (10 ng/mL) was used as a positive control (Biological replicate: N=24). Data are presented as mean±SEM. (C-G) Cells were treated with vehicle, dopamine, or a positive control (LPS, 10 ng/mL) in a time-course experiment (5, 15, 30, and 60 minutes). The phosphorylation ratios of AKT (Biological replicate: N=6-13), p38 (Biological replicate: N=6-11), p44/42 (Biological replicate: N=6), and GSK3β (Biological replicate: N=6-10) were analyzed by western blotting. Individual values are shown with line demonstrating mean. (H,I) hMDMs treated with dopamine for 30 minutes were assessed for the involvement of STAT signaling using spot blotting (Biological replicate: N=4). Matched individual values are shown with the bars demonstrating mean. (J,K) Cells were treated with vehicle or dopamine for 3 hrs, and gene expression levels of IFIT1 and ISG15, downstream targets of the IRF3 transcription factor, were assessed using qPCR (Biological replicate: N=7 for IFIT1 and N=5 for ISG15). Matched individual values are shown with the bar demonstrating mean. (L) Cells were treated with vehicle or dopamine for 6 hrs and cell counts were measured via CX7 imaging. (Biological replicate: N=6). Matched individual values are shown with the bar demonstrating mean. Paired comparisons were analyzed using a paired *t*-test (non-parametric: Wilcoxon test) in panel B. Significance levels are indicated as follows: *p* < 0.05 (*).

**Supplementary Figure 4.**
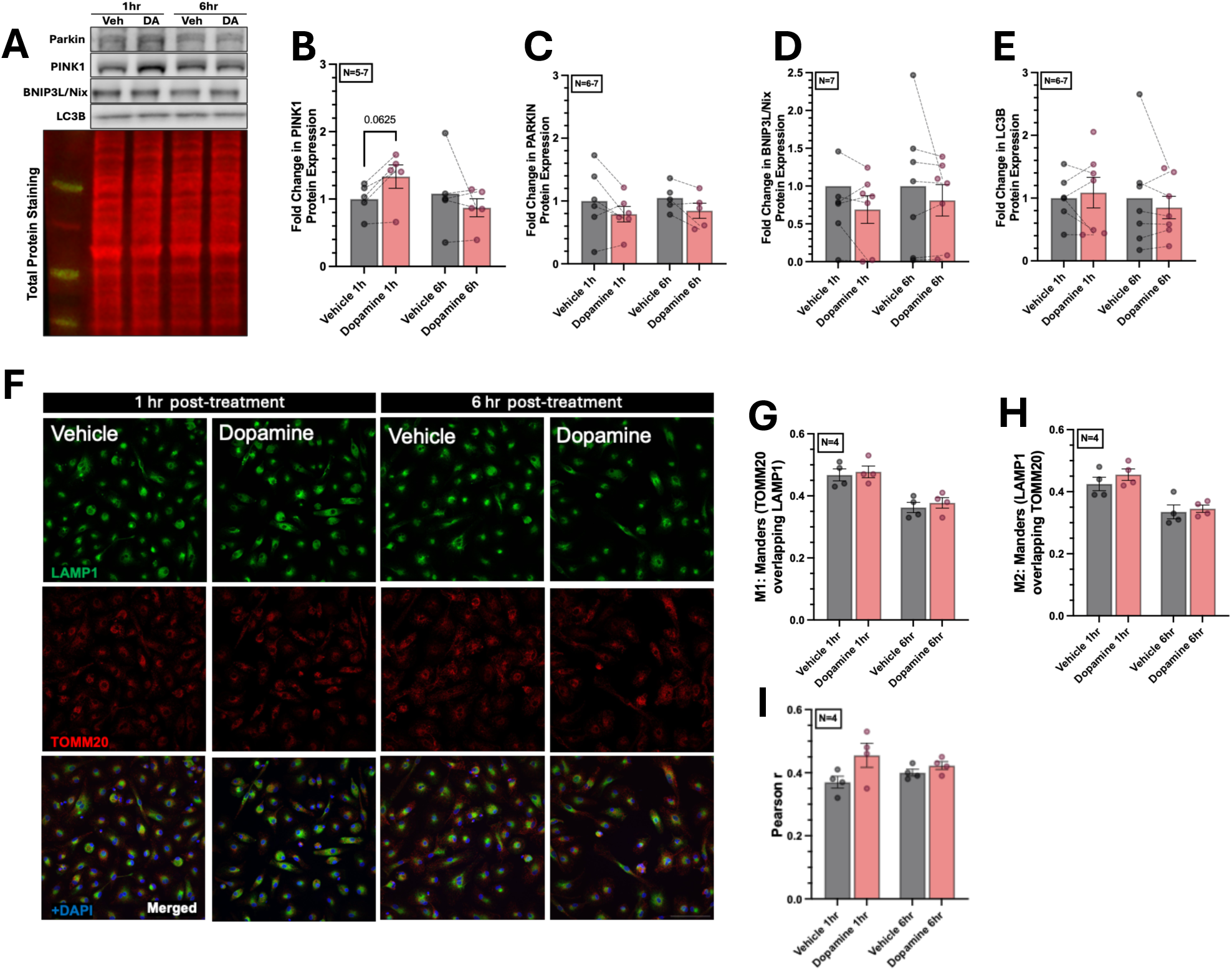
Dopamine does not induce mitophagy despite mitochondrial damage. (A-E) hMDMs were treated with dopamine and lysates were collected at 1hr or 6hr post-treatment, and were assessed for the expression levels of proteins involved in the mitophagy pathway, including PINK1 (Biological replicate: N=5-7), PARKIN (Biological replicate: N=6-7), BNIP3L/Nix (Biological replicate: N=7) and LC3B (Biological replicate: N=6-7) by western blotting. Data are presented as mean±SEM. Matched individual values are shown with the bar demonstrating mean. (F) Cells were stained with DAPI (blue), LAMP1 (green) and TOMM20 (red) to detect colocalization between lysosome and mitochondria to assess dopamine induced mitophagy overtime. cells were imaged using confocal microscopy (30× objective) with 3D acquisition at a step size of 0.15 μm. The scale bar represents 20 μm. Images were processed using ImageJ, and representative images are shown as z-projections. (G-I) Colocalization between LAMP1 and TOMM20 was quantified using the Coloc2 plugin in Fiji (ImageJ). Pearson’s correlation coefficient (R) and Manders’ overlap coefficients (M1 and M2) were calculated. (Biological replicate: N=4). Data are presented as mean±SEM. Paired comparisons were analyzed using paired one-way ANOVA (non-parametric: Friedman test) in panel B.

**Table 1.** Dopamine induces a pro-inflammatory cytokine profile in human macrophages revealed by MSD V-PLEX Analysis: hMDMs from three distinct donor groups were treated with vehicle or dopamine (1 μM) and collected 24 hrs post-stimulation. The table summarizes the relative cytokine production under each condition. ND indicates values below the limit of detection.

| Calculated Cytokine Concentration | Donor #1 |  | Donor #2 |  | Donor #3 |  |
| --- | --- | --- | --- | --- | --- | --- |
|  | Vehicle | Dopamine | Vehicle | Dopamine | Vehicle | Dopamine |
| **IL-6** | 23.824 | 138.376 | 158.860 | 239.930 | 131.399 | 154.823 |
| **IL-7** | 0.109 | 0.247 | 0.140 | 0.245 | ND | 0.168 |
| **IL-12p70** | 251.537 | ND | 273.366 | 275.120 | 295.627 | ND |
| **IL-2** | 1.974 | 5.550 | 5.335 | 9.788 | 10.408 | 5.486 |
| **IL-4** | 0.756 | 11.233 | 1.499 | 0.424 | 0.736 | 15.364 |
| **IL-17** | 33.331 | 136.620 | 26.105 | 41.352 | 26.105 | 41.352 |
| **IL-13** | 4.498 | 69.678 | 5.360 | 9.113 | 11.747 | 87.410 |
| **IFNγ** | 13.311 | 99.356 | 133.859 | 358.095 | 44.103 | 6.614 |
| **IL-8** | 1780.121 | 1707.058 | 1846.474 | 1898.137 | 1884.307 | 1761.097 |
| **IL-10** | 2.399 | 353.466 | 23.930 | 17.561 | 749.225 | 433.853 |
| **TNFα** | 3.277 | 8.834 | 21.814 | 38.349 | 103.576 | 47.379 |
| **GM-CSF** | 0.358 | 3.272 | 202.649 | 257.874 | 1.534 | 0.089 |
| **IL-5** | ND | ND | 5.754 | 16.336 | ND | ND |
| **IL-12** | 1.865 | 1.801 | 3.423 | 3.941 | 700.180 | 3.022 |
| **IL-15** | 0.435 | 0.474 | 1.018 | 1.236 | 0.604 | 0.215 |
| **TNFβ** | 0.468 | 0.380 | 6.362 | 12.240 | 7.602 | 0.475 |
| **VEGF-A** | 0.703 | 0.921 | ND | 0.171 | 0.305 | 9.938 |
| **Eotaxin-3** | 10.197 | 27.178 | 33.654 | 43.040 | 39.265 | 26.253 |
| **Eotaxin** | 4.035 | 13.088 | 8.807 | 8.295 | 16.885 | 21.644 |
| **CXCL10** | 1123.703 | 1351.268 | 3419.750 | 3233.901 | 6466.093 | 1129.146 |
| **MCP-1** | 5763.222 | 5237.748 | 5944.821 | 5401.527 | 6081.868 | 5265.108 |
| **MCP-4** | 356.837 | 322.712 | 1724.531 | 2119.716 | 1042.824 | 1040.402 |
| **MDC** | 3393.702 | 3277.259 | 75150.848 | 74195.719 | 73081.853 | 10741.640 |
| **MiP-1α** | 43.879 | 67.575 | 625.430 | 1977.975 | 3097.240 | 117.154 |
| **MiP-1β** | 112.663 | 132.542 | 1099.389 | 1830.356 | 4170.198 | 504.904 |
| **TARC** | 31.750 | 449.256 | 187.230 | 189.642 | 136.322 | 196.844 |
| **bFGF** | 47.169 | 55.695 | 29.227 | 24.871 | 36.041 | 34.922 |
| **Flt-1** | 2202.835 | 2032.100 | 16197.435 | 23648.516 | 23612.480 | 1666.021 |
| **PIGF** | 13.769 | 41.632 | 18.804 | ND | ND | ND |
| **Tie-2** | 362.769 | ND | 172.395 | ND | ND | ND |
| **VEGF-A** | ND | ND | 143.962 | 321.230 | ND | ND |
| **VEGF-C** | ND | ND | ND | ND | ND | ND |
| **VEGF-D** | 6294.178 | 5444.246 | 5706.979 | 5854.997 | 2825.613 | 2598.517 |

**Supplementary Table 1.** High Content Imaging NFkB Translocation Analysis. Table summarizes the parameters used for the NFkB nuclear translocation assay on CX7 high content imaging analysis software.

| **Condition** | **Value** |
| --- | --- |
| Ch1: Smooth Factor | 1 |
| Ch1: Thresholding (Fixed) | 500-700 |
| BackgroundCorrectionCh1 | 50 |
| Object.Ch1.Average Intensity.Ch1 | 0-6000 |
| Ch2: Smooth Factor | 3 |
| Ch2: Thresholding (Fixed) | 400-600 |
| BackgroundCorrectionCh2 | -255 |
| Object.Ch2.Average Intensity.Ch2 | -300 |
| ObjectAreaCh2 | 48.2-984.5 |
| Object.Average Intensity.Ch2 | 0-3701.25 |
| Ch3: Smooth Factor | 2 |
| Ch3: Thresholding (Fixed) | 400 |
| BackgroundCorrectionCh3 | -255 |
| Object.Ch3.Average Intensity.Ch3 | -3000 |
| ObjectAreaCh3 | 42.1-5826.16 |
| Object.Average Intensity.Ch3 | 0-4318 |
| ROI.A.Mask Ch | Channel 1 (DAPI) |
| ROI.A.Target_1 | Channel 2 (NF-kB) |
| ROI.B.Mask Ch | Channel 3 (CMDR) |
| ROI.B.Target_1 | Channel 2 (NF-kB) |
| ROI.B.Exclude | Channel 1 (DAPI) |
| RejectBorderObjects | Y |

**Supplementary Table 2.**
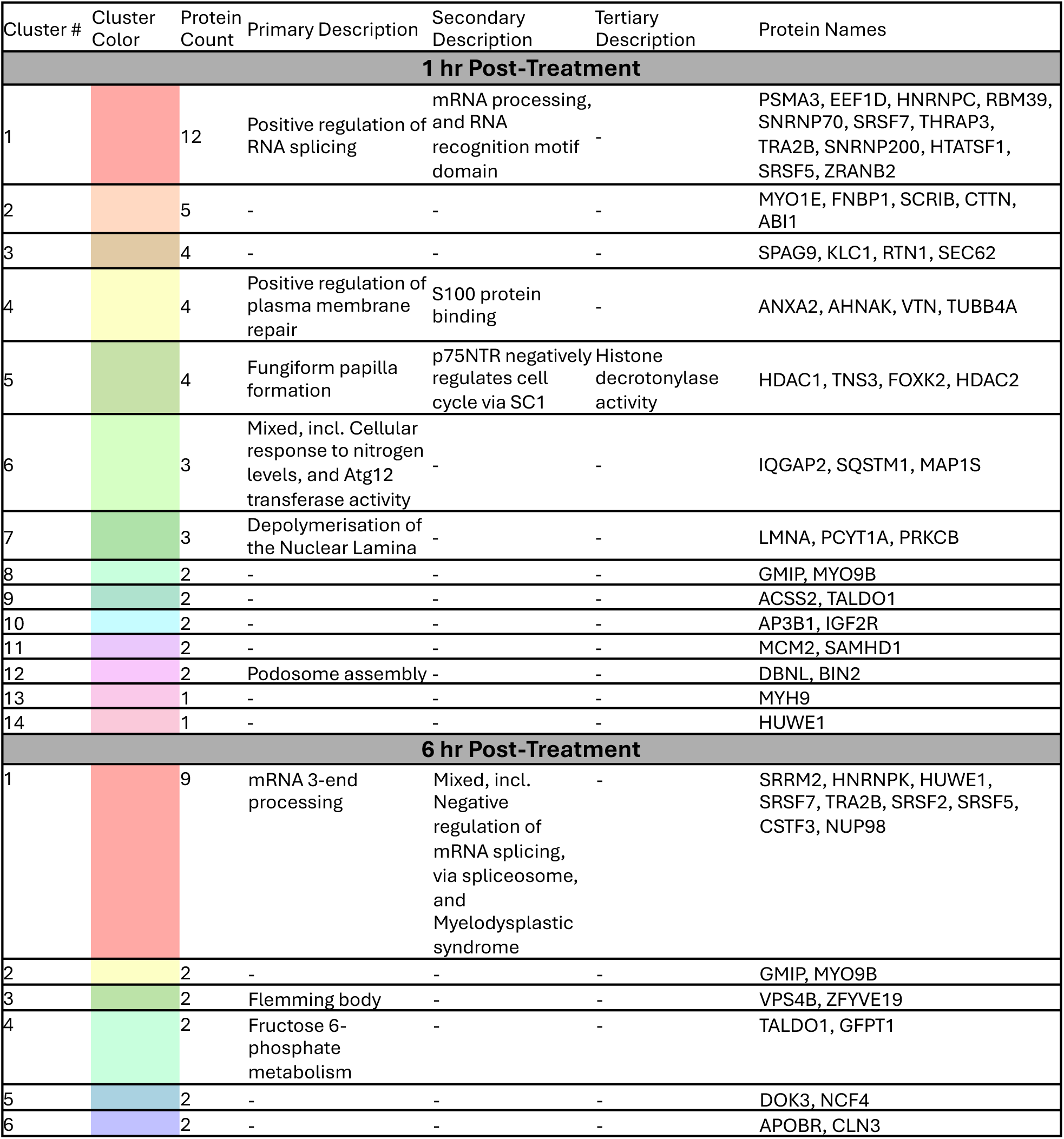
STRING network clustering shows time-dependent functional phospho-protein modules. Clusters of STRING biologically connected phospho-sites with ≥2-fold increased phosphorylation was performed via Markov Cluster (MCL) algorithm as shown in Supplementary Figure 2C,D and clusters are presented here. For each cluster, the table lists cluster number, color designation, protein count, and functional annotations, including primary, secondary, and tertiary descriptions derived from pathway and gene ontology enrichment analyses.

## References

1. Schultz, W. (2007). Multiple dopamine functions at different time courses. Annu Rev Neurosci 30, 259–288. 10.1146/annurev.neuro.28.061604.135722.

2. Schultz, W. (2016). Dopamine reward prediction-error signalling: a two-component response. Nat Rev Neurosci 17, 183–195. 10.1038/nrn.2015.26.

3. Kopec, A.M., Smith, C.J., and Bilbo, S.D. (2019). Neuro-Immune Mechanisms Regulating Social Behavior: Dopamine as Mediator? Trends Neurosci 42, 337–348. 10.1016/j.tins.2019.02.005.

4. Channer, B., Matt, S.M., Nickoloff-Bybel, E.A., Pappa, V., Agarwal, Y., Wickman, J., and Gaskill, P.J. (2023). Dopamine, Immunity, and Disease. Pharmacol Rev 75, 62–158. 10.1124/pharmrev.122.000618.

5. Gaskill, P.J., and Khoshbouei, H. (2022). Dopamine and norepinephrine are embracing their immune side and so should we. Curr Opin Neurobiol 77, 102626. 10.1016/j.conb.2022.102626.

6. Matt, S.M., and Gaskill, P.J. (2020). Where Is Dopamine and how do Immune Cells See it?: Dopamine-Mediated Immune Cell Function in Health and Disease. J Neuroimmune Pharmacol 15, 114–164. 10.1007/s11481-019-09851-4.

7. Huang, Y., Qiu, A.-W., Peng, Y.-P., Liu, Y., Huang, H.-W., and Qiu, Y.-H. (2010). Roles of dopamine receptor subtypes in mediating modulation of T lymphocyte function. Neuro Endocrinol Lett 31, 782–791.

8. Dobrodeeva, L.K., Samodova, A.V., Patrakeeva, V.P., Balashova, S.N., and Strekalovskaya, M.Yu. (2024). Increased Concentrations of Dopamine in the Blood and the State of the Immune System in Practically Healthy Residents of the Northern Territories. Hum Physiol 50, 521–528. 10.1134/S0362119724700993.

9. Matt, S.M., Nolan, R., Manikandan, S., Agarwal, Y., Channer, B., Oteju, O., Daniali, M., Canagarajah, J.A., LuPone, T., Mompho, K., et al. (2025). Dopamine-driven increase in IL-1β in myeloid cells is mediated by differential dopamine receptor expression and exacerbated by HIV. J Neuroinflammation 22, 91. 10.1186/s12974-025-03403-9.

10. Nolan, R.A., Reeb, K.L., Rong, Y., Matt, S.M., Johnson, H.S., Runner, K., and Gaskill, P.J. (2020). Dopamine activates NF-κB and primes the NLRP3 inflammasome in primary human macrophages. Brain Behav Immun Health 2, 100030. 10.1016/j.bbih.2019.100030.

11. Gopinath, A., Mackie, P.M., Phan, L.T., Mirabel, R., Smith, A.R., Miller, E., Franks, S., Syed, O., Riaz, T., Law, B.K., et al. (2023). Who Knew? Dopamine Transporter Activity Is Critical in Innate and Adaptive Immune Responses. Cells 12, 269. 10.3390/cells12020269.

12. Wieber, K., Fleige, L., Tsiami, S., Reinders, J., Braun, J., Baraliakos, X., and Capellino, S. (2022). Dopamine receptor 1 expressing B cells exert a proinflammatory role in female patients with rheumatoid arthritis. Sci Rep 12, 5985. 10.1038/s41598-022-09891-6.

13. Liu, X., Fan, T., Guan, J., Luo, A., Yu, Y., Chen, D., Mao, B., Jiang, H., and Liu, W. (2023). Dopamine relieves inflammatory responses through the D2 receptor after electroacupuncture at ST36 in a mouse model of chronic obstructive pulmonary disease. Acupunct Med 41, 163–174. 10.1177/09645284221107684.

14. Feng, Y., and Lu, Y. (2021). Immunomodulatory Effects of Dopamine in Inflammatory Diseases. Front. Immunol. 12, 663102. 10.3389/fimmu.2021.663102.

15. Gaskill, P.J., Yano, H.H., Kalpana, G.V., Javitch, J.A., and Berman, J.W. (2014). Dopamine Receptor Activation Increases HIV Entry into Primary Human Macrophages. PLoS ONE 9, e108232. 10.1371/journal.pone.0108232.

16. McKenna, F., McLaughlin, P.J., Lewis, B.J., Sibbring, G.C., Cummerson, J.A., Bowen-Jones, D., and Moots, R.J. (2002). Dopamine receptor expression on human T- and B-lymphocytes, monocytes, neutrophils, eosinophils and NK cells: a flow cytometric study. J Neuroimmunol 132, 34–40. 10.1016/s0165-5728(02)00280-1.

17. Nolan, R.A., Muir, R., Runner, K., Haddad, E.K., and Gaskill, P.J. (2019). Role of Macrophage Dopamine Receptors in Mediating Cytokine Production: Implications for Neuroinflammation in the Context of HIV-Associated Neurocognitive Disorders. J Neuroimmune Pharmacol 14, 134–156. 10.1007/s11481-018-9825-2.

18. Kawano, M., Takagi, R., Saika, K., Matsui, M., and Matsushita, S. (2018). Dopamine regulates cytokine secretion during innate and adaptive immune responses. Int Immunol 30, 591–606. 10.1093/intimm/dxy057.

19. Dorronsoro, A., Lang, V., Ferrin, I., Fernández-Rueda, J., Zabaleta, L., Pérez-Ruiz, E., Sepúlveda, P., and Trigueros, C. (2020). Intracellular role of IL-6 in mesenchymal stromal cell immunosuppression and proliferation. Sci Rep 10, 21853. 10.1038/s41598-020-78864-4.

20. Murakami, M., Kamimura, D., and Hirano, T. (2019). Pleiotropy and Specificity: Insights from the Interleukin 6 Family of Cytokines. Immunity 50, 812–831. 10.1016/j.immuni.2019.03.027.

21. Choy, E.H., De Benedetti, F., Takeuchi, T., Hashizume, M., John, M.R., and Kishimoto, T. (2020). Translating IL-6 biology into effective treatments. Nat Rev Rheumatol 16, 335–345. 10.1038/s41584-020-0419-z.

22. Weinberg, S.E., Sena, L.A., and Chandel, N.S. (2015). Mitochondria in the Regulation of Innate and Adaptive Immunity. Immunity 42, 406–417. 10.1016/j.immuni.2015.02.002.

23. Marques, E., Kramer, R., and Ryan, D.G. (2024). Multifaceted mitochondria in innate immunity. npj Metab Health Dis 2, 6. 10.1038/s44324-024-00008-3.

24. Hoffmann, R.F., Jonker, M.R., Brandenburg, S.M., De Bruin, H.G., Ten Hacken, N.H.T., Van Oosterhout, A.J.M., and Heijink, I.H. (2019). Mitochondrial dysfunction increases pro-inflammatory cytokine production and impairs repair and corticosteroid responsiveness in lung epithelium. Sci Rep 9, 15047. 10.1038/s41598-019-51517-x.

25. Xu, J., Wakai, M., Xiong, K., Yang, Y., Prabakaran, A., Wu, S., Ahrens, D., Molina-Portela, M.D.P., Ni, M., Bai, Y., et al. (2025). The pro-inflammatory cytokine IL6 suppresses mitochondrial function via the gp130-JAK1/STAT1/3-HIF1α/ERRα axis. Cell Reports 44, 115403. 10.1016/j.celrep.2025.115403.

26. Casey, A.M., Ryan, D.G., Prag, H.A., Chowdhury, S.R., Marques, E., Turner, K., Gruszczyk, A.V., Yang, M., Wolf, D.M., Miljkovic, J.Lj., et al. (2025). Pro-inflammatory macrophages produce mitochondria-derived superoxide by reverse electron transport at complex I that regulates IL-1β release during NLRP3 inflammasome activation. Nat Metab 7, 493–507. 10.1038/s42255-025-01224-x.

27. Wang, Y., Li, N., Zhang, X., and Horng, T. (2021). Mitochondrial metabolism regulates macrophage biology. J Biol Chem 297, 100904. 10.1016/j.jbc.2021.100904.

28. Channer, B., Daniali, M., Sheldon, L., Emanuel, K., Agarwal, Y., Kist, T., Murphy, B.J., Niu, M., Dampier, W., Fox, H., et al. (2025). Microenvironmental conditions and serum availability alter primary human macrophage NF-κB inflammatory response and function. J Leukoc Biol 117, qiaf071. 10.1093/jleuko/qiaf071.

29. Demichev, V., Messner, C.B., Vernardis, S.I., Lilley, K.S., and Ralser, M. (2020). DIA-NN: neural networks and interference correction enable deep proteome coverage in high throughput. Nat Methods 17, 41–44. 10.1038/s41592-019-0638-x.

30. Willforss, J., Chawade, A., and Levander, F. (2019). NormalyzerDE: Online Tool for Improved Normalization of Omics Expression Data and High-Sensitivity Differential Expression Analysis. J Proteome Res 18, 732–740. 10.1021/acs.jproteome.8b00523.

31. Shah, A.D., Goode, R.J.A., Huang, C., Powell, D.R., and Schittenhelm, R.B. (2020). LFQ-Analyst: An Easy-To-Use Interactive Web Platform To Analyze and Visualize Label-Free Proteomics Data Preprocessed with MaxQuant. J Proteome Res 19, 204–211. 10.1021/acs.jproteome.9b00496.

32. Chaudhry, A., Shi, R., and Luciani, D.S. (2020). A pipeline for multidimensional confocal analysis of mitochondrial morphology, function, and dynamics in pancreatic β-cells. Am J Physiol Endocrinol Metab 318, E87–E101. 10.1152/ajpendo.00457.2019.

33. Jiang, C.Y., Zhao, L., Green, M.D., Ravishankar, S., Towlerton, A.M.H., Scott, A.J., Raghavan, M., Cusick, M.F., Warren, E.H., and Ramnath, N. (2024). Class II HLA-DRB4 is a predictive biomarker for survival following immunotherapy in metastatic non-small cell lung cancer. Sci Rep 14, 345. 10.1038/s41598-023-48546-y.

34. Almeida, N., Carrara, G., Palmeira, C.M., Fernandes, A.S., Parsons, M., Smith, G.L., and Saraiva, N. (2020). Stimulation of cell invasion by the Golgi Ion Channel GAAP/TMBIM4 via an H2O2-Dependent Mechanism. Redox Biol 28, 101361. 10.1016/j.redox.2019.101361.

35. Pu, Y., Yang, J., Pan, Q., Li, C., Wang, L., Xie, X., Chen, X., Xiao, F., and Chen, G. (2024). MGST3 regulates BACE1 protein translation and amyloidogenesis by controlling the RGS4-mediated AKT signaling pathway. Journal of Biological Chemistry 300, 107530. 10.1016/j.jbc.2024.107530.

36. Adams, M.N., Croft, L.V., Urquhart, A., Saleem, M.A.M., Rockstroh, A., Duijf, P.H.G., Thomas, P.B., Ferguson, G.P., Najib, I.M., Shah, E.T., et al. (2023). hSSB1 (NABP2/OBFC2B) modulates the DNA damage and androgen-induced transcriptional response in prostate cancer. Prostate 83, 628–640. 10.1002/pros.24496.

37. Kolb, A., Kulis-Mandic, A.-M., Klein, M., Stastny, A., Haist, M., Votteler, V., Weidenthaler-Barth, B., Sinnberg, T., Sucker, A., Allies, G., et al. (2025). Constitutive expression of the transcriptional co-activator IκBζ promotes melanoma growth and immunotherapy resistance. Nat Commun 16, 5387. 10.1038/s41467-025-60929-5.

38. Jin, S., Chin, J., Seeber, S., Niewoehner, J., Weiser, B., Beaucamp, N., Woods, J., Murphy, C., Fanning, A., Shanahan, F., et al. (2013). TL1A/TNFSF15 directly induces proinflammatory cytokines, including TNFα, from CD3+CD161+ T cells to exacerbate gut inflammation. Mucosal Immunol 6, 886–899. 10.1038/mi.2012.124.

39. Guo, Q., Jin, Y., Chen, X., Ye, X., Shen, X., Lin, M., Zeng, C., Zhou, T., and Zhang, J. (2024). NF-κB in biology and targeted therapy: new insights and translational implications. Sig Transduct Target Ther 9, 53. 10.1038/s41392-024-01757-9.

40. Liu, T., Zhang, L., Joo, D., and Sun, S.-C. (2017). NF-κB signaling in inflammation. Sig Transduct Target Ther 2, 17023. 10.1038/sigtrans.2017.23.

41. Smith, J.A. (2020). STING, the Endoplasmic Reticulum, and Mitochondria: Is Three a Crowd or a Conversation? Front Immunol 11, 611347. 10.3389/fimmu.2020.611347.

42. Kim, J., Kim, H.-S., and Chung, J.H. (2023). Molecular mechanisms of mitochondrial DNA release and activation of the cGAS-STING pathway. Exp Mol Med 55, 510–519. 10.1038/s12276-023-00965-7.

43. Maekawa, H., Inoue, T., Ouchi, H., Jao, T.-M., Inoue, R., Nishi, H., Fujii, R., Ishidate, F., Tanaka, T., Tanaka, Y., et al. (2019). Mitochondrial Damage Causes Inflammation via cGAS-STING Signaling in Acute Kidney Injury. Cell Reports 29, 1261–1273.e6. 10.1016/j.celrep.2019.09.050.

44. Yum, S., Li, M., Fang, Y., and Chen, Z.J. (2021). TBK1 recruitment to STING activates both IRF3 and NF-κB that mediate immune defense against tumors and viral infections. Proc. Natl. Acad. Sci. U.S.A. 118, e2100225118. 10.1073/pnas.2100225118.

45. Zhang, L., Wei, X., Wang, Z., Liu, P., Hou, Y., Xu, Y., Su, H., Koci, M.D., Yin, H., and Zhang, C. (2023). NF-κB activation enhances STING signaling by altering microtubule-mediated STING trafficking. Cell Reports 42, 112185. 10.1016/j.celrep.2023.112185.

46. Li, B., Zhang, C., Xu, S., Li, Y., Vela, D., Vasquez, H., Zhang, L., Chakraborty, A., Lu, H.S., Coselli, J.S., et al. (2026). Epigenetic Programming of Macrophage Phenotypes by STING-IRF3 Drives Inflammation in Ascending Thoracic Aortic Dissection. Preprint at Immunology, 10.64898/2026.01.22.701198 https://doi.org/10.64898/2026.01.22.701198.

47. Basit, A., Cho, M.-G., Kim, E.-Y., Kwon, D., Kang, S.-J., and Lee, J.-H. (2020). The cGAS/STING/TBK1/IRF3 innate immunity pathway maintains chromosomal stability through regulation of p21 levels. Exp Mol Med 52, 643–657. 10.1038/s12276-020-0416-y.

48. Lama, L., Adura, C., Xie, W., Tomita, D., Kamei, T., Kuryavyi, V., Gogakos, T., Steinberg, J.I., Miller, M., Ramos-Espiritu, L., et al. (2019). Development of human cGAS-specific small-molecule inhibitors for repression of dsDNA-triggered interferon expression. Nat Commun 10, 2261. 10.1038/s41467-019-08620-4.

49. Hosoki, K., Govindhan, A., and Sur, S. (2025). Inhibition of STING by H151 attenuates Th2 differentiation and allergic lung inflammation. Journal of Allergy and Clinical Immunology 155, AB268. 10.1016/j.jaci.2024.12.829.

50. Szabo, M., Lagos, D., Cross, E., Collier, J., and Horvath, R. (2026). Mitochondrial DNA release and inflammation in mitochondrial disease pathogenesis. Brain, awag037. 10.1093/brain/awag037.

51. Kim, J., Gupta, R., Blanco, L.P., Yang, S., Shteinfer-Kuzmine, A., Wang, K., Zhu, J., Yoon, H.E., Wang, X., Kerkhofs, M., et al. (2019). VDAC oligomers form mitochondrial pores to release mtDNA fragments and promote lupus-like disease. Science 366, 1531–1536. 10.1126/science.aav4011.

52. Widdrington, J.D., Gomez-Duran, A., Steyn, J.S., Pyle, A., Ruchaud-Sparagano, M.-H., Scott, J., Baudouin, S.V., Rostron, A.J., Simpson, J., and Chinnery, P.F. (2017). Mitochondrial DNA depletion induces innate immune dysfunction rescued by IFN-γ. Journal of Allergy and Clinical Immunology 140, 1461–1464.e8. 10.1016/j.jaci.2017.04.048.

53. Lee, W., Choi, H.-I., Kim, M.-J., and Park, S.-Y. (2008). Depletion of mitochondrial DNA up-regulates the expression of MDR1 gene via an increase in mRNA stability. Exp Mol Med 40, 109. 10.3858/emm.2008.40.1.109.

54. Cierco, C., Santos, F., Nóbrega-Pereira, S., Da Cruz E Silva, O.A.B., and Trigo, D. (2026). Beyond Pulsing Dyes: Are Flickers the Language of the Mitochondrial Network? Preprint at Cell Biology, 10.64898/2026.03.24.713912 https://doi.org/10.64898/2026.03.24.713912.

55. Pouvreau, S. (2010). Mitochondrial Superoxide Flashes in Skeletal Muscle are Linked to Metabolism. Biophysical Journal 98, 377a. 10.1016/j.bpj.2009.12.2033.

56. Lin, A.J., Joshi, A.U., Mukherjee, R., Tompkins, C.A., Vijayan, V., Mochly-Rosen, D., and Haileselassie, B. (2022). δPKC-Mediated DRP1 Phosphorylation Impacts Macrophage Mitochondrial Function and Inflammatory Response to Endotoxin. Shock 57, 435–443. 10.1097/SHK.0000000000001885.

57. Lin, Y., Wang, D., Li, B., Wang, J., Xu, L., Sun, X., Ji, K., Yan, C., Liu, F., and Zhao, Y. (2024). Targeting DRP1 with Mdivi-1 to correct mitochondrial abnormalities in ADOA+ syndrome. JCI Insight 9, e180582. 10.1172/jci.insight.180582.

58. Bordt, E.A., Clerc, P., Roelofs, B.A., Saladino, A.J., Tretter, L., Adam-Vizi, V., Cherok, E., Khalil, A., Yadava, N., Ge, S.X., et al. (2017). The Putative Drp1 Inhibitor mdivi-1 Is a Reversible Mitochondrial Complex I Inhibitor that Modulates Reactive Oxygen Species. Developmental Cell 40, 583–594.e6. 10.1016/j.devcel.2017.02.020.

59. Deragon, M.A., McCaig, W.D., Patel, P.S., Haluska, R.J., Hodges, A.L., Sosunov, S.A., Murphy, M.P., Ten, V.S., and LaRocca, T.J. (2020). Mitochondrial ROS prime the hyperglycemic shift from apoptosis to necroptosis. Cell Death Discov. 6, 132. 10.1038/s41420-020-00370-3.

60. Hao, X., Bu, W., Lv, G., Xu, L., Hou, D., Wang, J., Liu, X., Yang, T., Zhang, X., Liu, Q., et al. (2022). Disrupted mitochondrial homeostasis coupled with mitotic arrest generates antineoplastic oxidative stress. Oncogene 41, 427–443. 10.1038/s41388-021-02105-9.

61. Yu, Q., Wang, Y., Dong, L., He, Y., Liu, R., Yang, Q., Cao, Y., Wang, Y., Jia, A., Bi, Y., et al. (2020). Regulations of Glycolytic Activities on Macrophages Functions in Tumor and Infectious Inflammation. Front. Cell. Infect. Microbiol. 10, 287. 10.3389/fcimb.2020.00287.

62. Wculek, S.K., Dunphy, G., Heras-Murillo, I., Mastrangelo, A., and Sancho, D. (2022). Metabolism of tissue macrophages in homeostasis and pathology. Cell Mol Immunol 19, 384–408. 10.1038/s41423-021-00791-9.

63. Kim, I., Rodriguez-Enriquez, S., and Lemasters, J.J. (2007). Selective degradation of mitochondria by mitophagy. Archives of Biochemistry and Biophysics 462, 245–253. 10.1016/j.abb.2007.03.034.

64. Küng, C., Lazarou, M., and Nguyen, T.N. (2025). Advances in mitophagy initiation mechanisms. Current Opinion in Cell Biology 94, 102493. 10.1016/j.ceb.2025.102493.

65. Bhatia, D., Chung, K.-P., Nakahira, K., Patino, E., Rice, M.C., Torres, L.K., Muthukumar, T., Choi, A.M., Akchurin, O.M., and Choi, M.E. (2019). Mitophagy-dependent macrophage reprogramming protects against kidney fibrosis. JCI Insight 4, e132826, 132826. 10.1172/jci.insight.132826.

66. Ketchum, F., Celebi, L., Hawthorne, L., and Zorlutuna, P. (2025). Mitochondria Clearance Enables Macrophage-Driven Maturation of iPSC-Derived Cardiomyocyte Metabolism. Preprint at Bioengineering, 10.1101/2025.09.02.673264 https://doi.org/10.1101/2025.09.02.673264.

67. Choudhuri, S., Chowdhury, I.H., and Garg, N.J. (2020). Mitochondrial Regulation of Macrophage Response Against Pathogens. Front Immunol 11, 622602. 10.3389/fimmu.2020.622602.

68. Tannahill, G.M., Curtis, A.M., Adamik, J., Palsson-McDermott, E.M., McGettrick, A.F., Goel, G., Frezza, C., Bernard, N.J., Kelly, B., Foley, N.H., et al. (2013). Succinate is an inflammatory signal that induces IL-1β through HIF-1α. Nature 496, 238–242. 10.1038/nature11986.

69. O’Neill, L.A.J., and Pearce, E.J. (2016). Immunometabolism governs dendritic cell and macrophage function. J Exp Med 213, 15–23. 10.1084/jem.20151570.

70. Wong, K.Y., Roy, J., Fung, M.L., Heng, B.C., Zhang, C., and Lim, L.W. (2020). Relationships between Mitochondrial Dysfunction and Neurotransmission Failure in Alzheimer’s Disease. Aging Dis 11, 1291–1316. 10.14336/AD.2019.1125.

71. Rawani, N.S., Chan, A.W., Dursun, S.M., and Baker, G.B. (2024). The Underlying Neurobiological Mechanisms of Psychosis: Focus on Neurotransmission Dysregulation, Neuroinflammation, Oxidative Stress, and Mitochondrial Dysfunction. Antioxidants 13, 709. 10.3390/antiox13060709.

72. Sharma, N., Jain, M., and Navik, U. (2026). Mitochondrial Health and Critical Neurotransmitters in Neurodegeneration. In Neuroendocrine-Immune Axis, R. Shukla, S. Naqvi, M. S. Deore, and R. K. Kaundal, eds. (Springer Nature Singapore), pp. 247-273. 10.1007/978-981-95-5088-3_11.

73. Kuznetsov, A.V., Javadov, S., Saks, V., Margreiter, R., and Grimm, M. (2017). Synchronism in mitochondrial ROS flashes, membrane depolarization and calcium sparks in human carcinoma cells. Biochim Biophys Acta Bioenerg 1858, 418–431. 10.1016/j.bbabio.2017.03.001.

74. Narendra, D.P., and Youle, R.J. (2024). The role of PINK1-Parkin in mitochondrial quality control. Nat Cell Biol 26, 1639–1651. 10.1038/s41556-024-01513-9.

75. Luciani, A., Schumann, A., Berquez, M., Chen, Z., Nieri, D., Failli, M., Debaix, H., Festa, B.P., Tokonami, N., Raimondi, A., et al. (2020). Impaired mitophagy links mitochondrial disease to epithelial stress in methylmalonyl-CoA mutase deficiency. Nat Commun 11, 970. 10.1038/s41467-020-14729-8.

76. Rebetz, J., Cederholm, H., McGauran, D., Moore, E., Chi, C., Tabak, K., Allhorn, M., Olsson, M.L., Egesten, A., Semple, J.W., et al. (2025). Mitochondrial DNA via recipient TLR9 acts as a potent first hit in murine transfusion-related acute lung injury. Blood 146, 2479–2490. 10.1182/blood.2025028794.

77. Yang, N.-S.-Y., Zhong, W.-J., Sha, H.-X., Zhang, C.-Y., Jin, L., Duan, J.-X., Xiong, J.-B., You, Z.-J., Zhou, Y., and Guan, C.-X. (2024). mtDNA-cGAS-STING axis-dependent NLRP3 inflammasome activation contributes to postoperative cognitive dysfunction induced by sevoflurane in mice. Int J Biol Sci 20, 1927–1946. 10.7150/ijbs.91543.

78. Lei, Y., VanPortfliet, J.J., Chen, Y.-F., Bryant, J.D., Li, Y., Fails, D., Torres-Odio, S., Ragan, K.B., Deng, J., Mohan, A., et al. (2023). Cooperative sensing of mitochondrial DNA by ZBP1 and cGAS promotes cardiotoxicity. Cell 186, 3013–3032.e22. 10.1016/j.cell.2023.05.039.

79. Shimada, K., Crother, T.R., Karlin, J., Dagvadorj, J., Chiba, N., Chen, S., Ramanujan, V.K., Wolf, A.J., Vergnes, L., Ojcius, D.M., et al. (2012). Oxidized Mitochondrial DNA Activates the NLRP3 Inflammasome during Apoptosis. Immunity 36, 401–414. 10.1016/j.immuni.2012.01.009.

80. Xu, L., Zhou, J., Che, J., Wang, H., Yang, W., Zhou, W., and Zhao, H. (2021). Mitochondrial DNA enables AIM2 inflammasome activation and hepatocyte pyroptosis in nonalcoholic fatty liver disease. Am J Physiol Gastrointest Liver Physiol 320, G1034–G1044. 10.1152/ajpgi.00431.2020.

81. Perry, A.K., Chow, E.K., Goodnough, J.B., Yeh, W.-C., and Cheng, G. (2004). Differential requirement for TANK-binding kinase-1 in type I interferon responses to toll-like receptor activation and viral infection. J Exp Med 199, 1651–1658. 10.1084/jem.20040528.

82. Hemmi, H., Takeuchi, O., Sato, S., Yamamoto, M., Kaisho, T., Sanjo, H., Kawai, T., Hoshino, K., Takeda, K., and Akira, S. (2004). The roles of two IkappaB kinase-related kinases in lipopolysaccharide and double stranded RNA signaling and viral infection. J Exp Med 199, 1641–1650. 10.1084/jem.20040520.

83. Clark, K., Plater, L., Peggie, M., and Cohen, P. (2009). Use of the pharmacological inhibitor BX795 to study the regulation and physiological roles of TBK1 and IkappaB kinase epsilon: a distinct upstream kinase mediates Ser-172 phosphorylation and activation. J Biol Chem 284, 14136–14146. 10.1074/jbc.M109.000414.

84. Dunphy, G., Flannery, S.M., Almine, J.F., Connolly, D.J., Paulus, C., Jønsson, K.L., Jakobsen, M.R., Nevels, M.M., Bowie, A.G., and Unterholzner, L. (2018). Non-canonical Activation of the DNA Sensing Adaptor STING by ATM and IFI16 Mediates NF-κB Signaling after Nuclear DNA Damage. Mol Cell 71, 745–760.e5. 10.1016/j.molcel.2018.07.034.

85. Stempel, M., Chan, B., Juranić Lisnić, V., Krmpotić, A., Hartung, J., Paludan, S.R., Füllbrunn, N., Lemmermann, N.A., and Brinkmann, M.M. (2019). The herpesviral antagonist m152 reveals differential activation of STING-dependent IRF and NF-κB signaling and STING’s dual role during MCMV infection. EMBO J 38, e100983. 10.15252/embj.2018100983.

86. Gopinath, A., Badov, M., Francis, M., Shaw, G., Collins, A., Miller, D.R., Hansen, C.A., Mackie, P., Tansey, M.G., Dagra, A., et al. (2021). TNFα increases tyrosine hydroxylase expression in human monocytes. NPJ Parkinsons Dis 7, 62. 10.1038/s41531-021-00201-x.

87. Zaparte, A., Schuch, J.B., Viola, T.W., Baptista, T.A.S., Beidacki, A.S., Do Prado, C.H., Sanvicente-Vieira, B., Bauer, M.E., and Grassi-Oliveira, R. (2019). Cocaine Use Disorder Is Associated With Changes in Th1/Th2/Th17 Cytokines and Lymphocytes Subsets. Front. Immunol. 10, 2435. 10.3389/fimmu.2019.02435.

88. Mestas, J., and Hughes, C.C.W. (2004). Of mice and not men: differences between mouse and human immunology. J Immunol 172, 2731–2738. 10.4049/jimmunol.172.5.2731.

89. Takao, K., and Miyakawa, T. (2015). Genomic responses in mouse models greatly mimic human inflammatory diseases. Proc. Natl. Acad. Sci. U.S.A. 112, 1167–1172. 10.1073/pnas.1401965111.

90. Kodamullil, A.T., Iyappan, A., Karki, R., Madan, S., Younesi, E., and Hofmann-Apitius, M. (2017). Of Mice and Men: Comparative Analysis of Neuro-Inflammatory Mechanisms in Human and Mouse Using Cause-and-Effect Models. J Alzheimers Dis 59, 1045–1055. 10.3233/JAD-170255.

91. Araki, K.Y., Sims, J.R., and Bhide, P.G. (2007). Dopamine receptor mRNA and protein expression in the mouse corpus striatum and cerebral cortex during pre- and postnatal development. Brain Res 1156, 31–45. 10.1016/j.brainres.2007.04.043.

92. Bjerke, I.E., Cullity, E.R., Kjelsberg, K., Charan, K.M., Leergaard, T.B., and Kim, J.H. (2022). DOPAMAP, high-resolution images of dopamine 1 and 2 receptor expression in developing and adult mouse brains. Sci Data 9, 175. 10.1038/s41597-022-01268-8.

93. Hollon, T.R., Bek, M.J., Lachowicz, J.E., Ariano, M.A., Mezey, E., Ramachandran, R., Wersinger, S.R., Soares-da-Silva, P., Liu, Z.F., Grinberg, A., et al. (2002). Mice Lacking D_5_ Dopamine Receptors Have Increased Sympathetic Tone and Are Hypertensive. J. Neurosci. 22, 10801–10810. 10.1523/JNEUROSCI.22-24-10801.2002.

94. Osorio-Barrios, F., Navarro, G., Campos, J., Ugalde, V., Prado, C., Raïch, I., Contreras, F., López, E., Espinoza, A., Lladser, A., et al. (2021). The Heteromeric Complex Formed by Dopamine Receptor D5 and CCR9 Leads the Gut Homing of CD4+ T Cells Upon Inflammation. Cell Mol Gastroenterol Hepatol 12, 489–506. 10.1016/j.jcmgh.2021.04.006.

95. Morimoto, K., Ouchi, M., Kitano, T., Eguchi, R., and Otsuguro, K. (2022). Dopamine regulates astrocytic IL-6 expression and process formation via dopamine receptors and adrenoceptors. European Journal of Pharmacology 928, 175110. 10.1016/j.ejphar.2022.175110.

96. Zhang, M., Chen, T., Lu, X., Lan, X., Chen, Z., and Lu, S. (2024). G protein-coupled receptors (GPCRs): advances in structures, mechanisms and drug discovery. Sig Transduct Target Ther 9, 88. 10.1038/s41392-024-01803-6.

97. Zhuang, Y., Xu, P., Mao, C., Wang, L., Krumm, B., Zhou, X.E., Huang, S., Liu, H., Cheng, X., Huang, X.-P., et al. (2021). Structural insights into the human D1 and D2 dopamine receptor signaling complexes. Cell 184, 931–942.e18. 10.1016/j.cell.2021.01.027.

98. Li, Y., Zhang, Y., Zhang, C., Wang, H., Wei, X., Chen, P., and Lu, L. (2020). Mitochondrial dysfunctions trigger the calcium signaling-dependent fungal multidrug resistance. Proc. Natl. Acad. Sci. U.S.A. 117, 1711–1721. 10.1073/pnas.1911560116.

99. Seegren, P.V., Harper, L.R., Downs, T.K., Zhao, X.-Y., Viswanathan, S.B., Stremska, M.E., Olson, R.J., Kennedy, J., Ewald, S.E., Kumar, P., et al. (2023). Reduced mitochondrial calcium uptake in macrophages is a major driver of inflammaging. Nat Aging 3, 796–812. 10.1038/s43587-023-00436-8.

100. Valenzuela, R., Costa-Besada, M.A., Iglesias-Gonzalez, J., Perez-Costas, E., Villar-Cheda, B., Garrido-Gil, P., Melendez-Ferro, M., Soto-Otero, R., Lanciego, J.L., Henrion, D., et al. (2016). Mitochondrial angiotensin receptors in dopaminergic neurons. Role in cell protection and aging-related vulnerability to neurodegeneration. Cell Death Dis 7, e2427–e2427. 10.1038/cddis.2016.327.

101. Puri, N.M., Romano, G.R., Lin, T.-Y., Mai, Q.N., and Irannejad, R. (2022). The organic cation transporter 2 regulates dopamine D1 receptor signaling at the Golgi apparatus. Elife 11, e75468. 10.7554/eLife.75468.

102. Mackie, P.M., Gopinath, A., Montas, D.M., Nielsen, A., Smith, A., Nolan, R.A., Runner, K., Matt, S.M., McNamee, J., Riklan, J.E., et al. (2022). Functional characterization of the biogenic amine transporters on human macrophages. JCI Insight 7, e151892. 10.1172/jci.insight.151892.

